# Reinforcement targets sexual or postmating prezygotic reproductive barriers depending on species abundance and population history

**DOI:** 10.1101/425058

**Authors:** Noora Poikela, Johanna Kinnunen, Mareike Wurdack, Hannele Kauranen, Thomas Schmitt, Maaria Kankare, Rhonda R. Snook, Anneli Hoikkala

**Affiliations:** Department of Biological and Environmental Science, University of Jyväskylä, Finland; Department of Animal Ecology and Tropical Biology, University of Wuerzburg, Germany; Department of Zoology, Stockholm University, Sweden

**Author notes:** **Corresponding author:** Noora Poikela, Department of Biological and Environmental Science, P.O. Box 35, FI-40014 University of Jyväskylä, Finland.

**Keywords:** speciation, sympatry, allopatry, female discrimination, courtship cue, Drosophila

## Abstract

The impact of different reproductive barriers on species or population isolation may vary in different stages of speciation depending on evolutionary forces acting within species and through species’ interactions. Genetic incompatibilities between interacting species are expected to reinforce prezygotic barriers in sympatric populations and create character displacement between conspecific populations living within and outside the area of sympatry. The outcome of reinforcement has been suggested to be affected by the strength of postzygotic barriers, the history of species coexistence, and the impact of species abundancies on females’ discrimination against heterospecific males. We tested these predictions in *Drosophila montana* and *Drosophila flavomontana* populations from different geographic regimes. All barriers between *D. montana* females and *D. flavomontana* males were extremely strong, while in the reciprocal cross postzygotic isolation was less effective and the target of reinforcement varied according to population type. In long-established sympatric populations, where *D. flavomontana* is abundant, reinforcement targeted sexual isolation, and in populations, where this species is a new invader and rare, reinforcement targeted postmating prezygotic barriers. Reinforcement of these barriers also created respective barriers between different *D. flavomontana* populations. These findings show that interspecies interactions have far-reaching effects on strengthening species barriers and promoting speciation.

## Introduction

Past and present climate change and human activity have induced shifts in species’ distribution, which has had a strong impact on species interactions and speciation. When geographically or ecologically isolated populations or diverging species spread in the same area/habitat, their interaction may lead to different evolutionary outcomes depending on the strength of the reproductive barriers that they have evolved during isolation. If the barriers are weak to moderate, then the gene pools of the evolving species may be either merged (Servedio and Noor 2003; Arnold and Martin 2009) or the species may exchange gene alleles via hybridization and backcrossing (Abbott et al. 2013). If the barriers are strong enough, then the two species or isolated populations may live in sympatry. If the barriers are not complete, and maladaptive hybridization occurs, then selection for reinforcement of barriers that function at an earlier stage of interactions between heterospecific individuals is predicted (Dobzhansky 1940; Howard 1993; Servedio and Noor 2003; Turissini et al. 2018). This reinforcement of sexual or postmating prezygotic (PMPZ) barriers will lead to reproductive character displacement (i.e. greater divergence between species in areas of sympatry than in areas of allopatry) in traits like female mate discrimination and preferences, courtship cues and gamete recognition. As a consequence, speciation between populations of the same species may be promoted (Howard 1993; Ortiz-Barrientos et al. 2009; Hoskin and Higgie 2010). Thus, to understand how the evolution of different components of reproductive isolation during species divergence occurs, and its broader implications, elucidating which barriers are targeted by reinforcement and the role of reinforcement in completing or initiating speciation both between species and between populations of a species is critical (Butlin et al. 2008; Nosil et al. 2009; The Marie Curie speciation network 2012).

One group of organisms in which the evolution of reproductive barriers has been well-studied is Drosophila. Sexual isolation in this taxon has been shown to evolve faster than postzygotic isolation (Coyne and Orr 1997), and PMPZ isolation faster than hybrid inviability but more slowly than sexual isolation (Turissini et al. 2018). However, there is no general agreement on how strongly reinforcement contributes to the evolution of prezygotic barriers. Sexual isolation is usually maintained by females, based on species differences in male-female interactions, courtship cues and female discrimination for these cues (Chenoweth and Blows 2006). Female acceptance threshold may vary between the interacting species, and different sensory modalities and courtship cues used in courtship and mating may differ between closely-related species (Gleason et al. 2012; Giglio and Dyer 2013; Colyott et al. 2016). Reinforcement enhances female discrimination against heterospecific males in sympatric populations and increases their discrimination towards conspecific males, and thus sympatric females may reject allopatric males as mates. The converse (i.e., allopatric females with sympatric males) need not be true (Noor 1999; Hoskin et al. 2005; Jaenike et al. 2006; Bewick and Dyer 2014). Most of the identified PMPZ barriers, including incompatibilities in the transfer, storage and use of heterospecific sperm, involve discordant interactions between gametes or between the female reproductive tract and male seminal fluids, and they can function after a single mating (Howard 1999; Wirtz 1999; Price et al. 2001; Howard et al. 2009). Even though postmating interactions can have important fitness consequences for both sexes, reinforcement of PMPZ barriers in insect species has been reported only between *D. yakuba* and *D. santomea* (Matute 2010) and between *D. pseudoobscura* and *D. persimilis* (Castillo and Moyle 2017).

Reinforcement is most likely when species hybridization is common and its costs are high, and when the opposing forces of gene flow and recombination are weak (e.g. Servedio and Noor 2003; Coyne and Orr 2004; Servedio 2009; Butlin and Smadja 2018). Accordingly, almost all sympatric Drosophila species have been found to have concordant pre- and post- zygotic isolation asymmetries, where the more costly reciprocal mating has greater prezygotic isolation relative to the less costly mating, while no such patterns exist in allopatry (Yukilevich 2012). The outcome of reinforcement can also be affected by changes in species’ distribution and abundance, the length of species coexistence, and the effects of natural and sexual selection between and within species (Servedio 2001; Servedio and Noor 2003; Smadja and Butlin 2011; Nosil 2012). Whether this strengthens or weakens female discrimination of heterospecific males is less clear. In the “rarer female hypothesis”, species recognition ability of females of the less abundant species is expected to get reinforced, because these females encounter more heterotypic mating attempts in the wild and suffer from higher hybridization costs than those of the more abundant species (Noor 1995; Hoskin et al. 2005; Yukilevich 2012). However, several studies have shown that females’ ability to distinguish hetero- from con-specifics weakens when population density is small and the likelihood of encountering conspecific males is low (see Wirtz 1999 for a review; Matute 2014). Reinforcement of sexual isolation in this scenario may not be possible, and thus natural and sexual selection could drive the evolution of PMPZ barriers to limit costs of maladaptive hybridization (Turissini et al. 2018).

Despite these predictions, few studies have examined whether the targets and consequences of reinforcement vary between species in different contexts – between species that have a longer history of sympatry compared to a scenario in which one species has only recently invaded and is still rare – and whether this variation impacts reproductive barriers between populations of the same species. We use the species pair, *D. montana* and *D. flavomontana*, which offers an excellent opportunity to address these outstanding speciation questions. The species diverged from each other from ~1 million (Poikela and Lohse et al., unpublished data) to 4.9 million years ago (Morales-hojas et al. 2011), and chromosomal studies performed on these species suggest that both of them originated from the Rocky Mountains (Stone et al. 1960), where they still hybridize to some degree (Patterson 1952). *D. montana* has distributed around the northern hemisphere, including the western coast of North America (Throckmorton 1982), while *D. flavomontana* has spread from the Rocky Mountains to the western coast only after the extensive collections carried out on this area in 1950’s (see Patterson 1952), and is still rare. Both species have a patchy population structure, as they live only on the waterside and as their distribution and abundance depend on climatic factors and the presence of species-specific host trees (*D. montana* aspen and alder and *D. flavomontana* cotton wood; Patterson 1952). Reproductive barriers between *D. montana* females and *D. flavomontana* males are nearly complete, while the barriers between *D. flavomontana* females and *D. montana* males are weaker (Patterson 1952), which provides an opportunity for reinforcement and its potential effects on reproductive isolation between conspecific populations.

Using this system, we have (1) studied the strength of postzygotic barriers between *D. montana* and *D. flavomontana* in allopatric populations and in sympatric populations with different histories and species abundances, (2) tested whether and how the strength of postzygotic barriers, the length of species coexistence and the species’ relative abundances have affected the reinforcement of sexual and/or PMPZ barriers, and (3) traced the effects of reinforcement on the divergence of reproductive traits and the enhancement of reproductive barriers between *D. flavomontana* populations from allopatry and sympatry (Fig. 1). We predict that in sympatric Rocky Mountains populations, where *D. flavomontana* is abundant, reinforcement has increased the discrimination of *D. flavomontana* females against *D. montana* males, and induced changes in the key courtship cues of *D. flavomontana* males, which has generated sexual isolation between the females of these populations and conspecific males from other populations (Fig. 1). Reinforcement may have occurred similarly in sympatric western coast populations, where *D. flavomontana* is a new invader and still rare, if species recognition of females has not decreased due to the lack of conspecific mating partners. If, however, female *D. flavomontana* mate recognition is low, reinforcement should have targeted PMPZ barriers and induced reproductive character displacement in traits maintaining these barriers both between the species and between *D. flavomontana* populations (Fig. 1).

**Figure 1.**
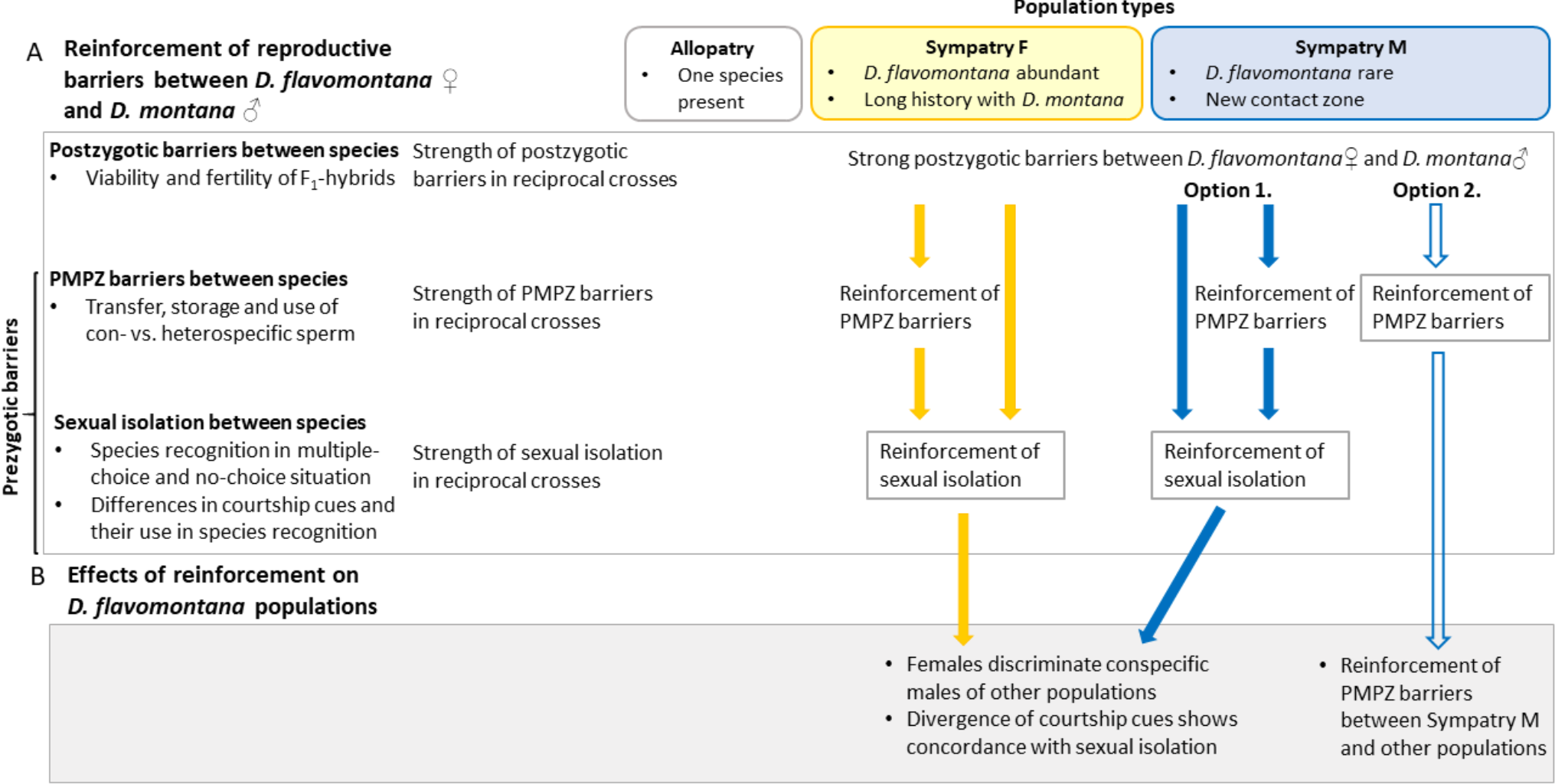
Predictions for the evolution of reproductive barriers. (A) The strength and asymmetry of reproductive barriers between *D. montana* and *D. flavomontana* in three population types, and reinforcement of prezygotic barriers in sympatric populations with different species abundances and different length of species coexistence (Sympatry F and Sympatry M). (B) Effects of reinforcement on divergence of reproductive traits among *D. flavomontana* populations.

## Material and Methods

### STUDY SPECIES

#### D. montana and D. flavomontana populations

*D. montana* and *D. flavomontana* belong to the montana subphylad of the *D. virilis* group (Morales-hojas et al. 2011). *D. montana* is distributed around the northern hemisphere and in North America it is found in high latitudes in Canada and Alaska, in high altitudes (from 1400 to above 3000 m) and wide range of latitudes in the Rocky Mountains, and in low altitudes and latitudes along the western coast of the United States (US) and Canadian Pacific coast (Patterson 1952; Stone et al. 1960; Throckmorton 1982). *D. flavomontana* lives in lower altitudes than *D. montana* (usually below 2000 m), and in the 1950s its distribution was restricted to the Rocky Mountains area (Patterson 1952; Stone et al. 1960). Our collections in 2010 – 2015 showed that the distribution of both species had shifted northwards and towards higher altitudes and that *D. flavomontana* had invaded the North American western coast, where it had not been detected before.

The allopatric strains used in this study are either truly allopatric (*D. montana*, Seward, Alaska) or from single-species sites on the Rocky Mountains area (*D. montana:* Afton, Wyoming; *D*. *flavomontana*: Livingston, Montana and Liberty, Utah; Fig. 2). Two types of sympatric strains were studied: collections representing the old distribution area of both species around the lower slopes of the Rocky Mountains (altitude up to 2 000 meters), where *D. flavomontana* is more abundant than *D. montana* (hereafter referred to as “Sympatry F”; Cranbrook, Canada and Jackson, Wyoming), and those from the western coast of North America, where *D. flavomontana* has invaded recently and is still rare (hereafter referred to as “Sympatry M”; Terrace, Canada; Vancouver, Canada; Ashford Washington). In the following, we refer to the origin of the strains (Allopatry, Sympatry F and M) as “population type”.

**Figure 2.**
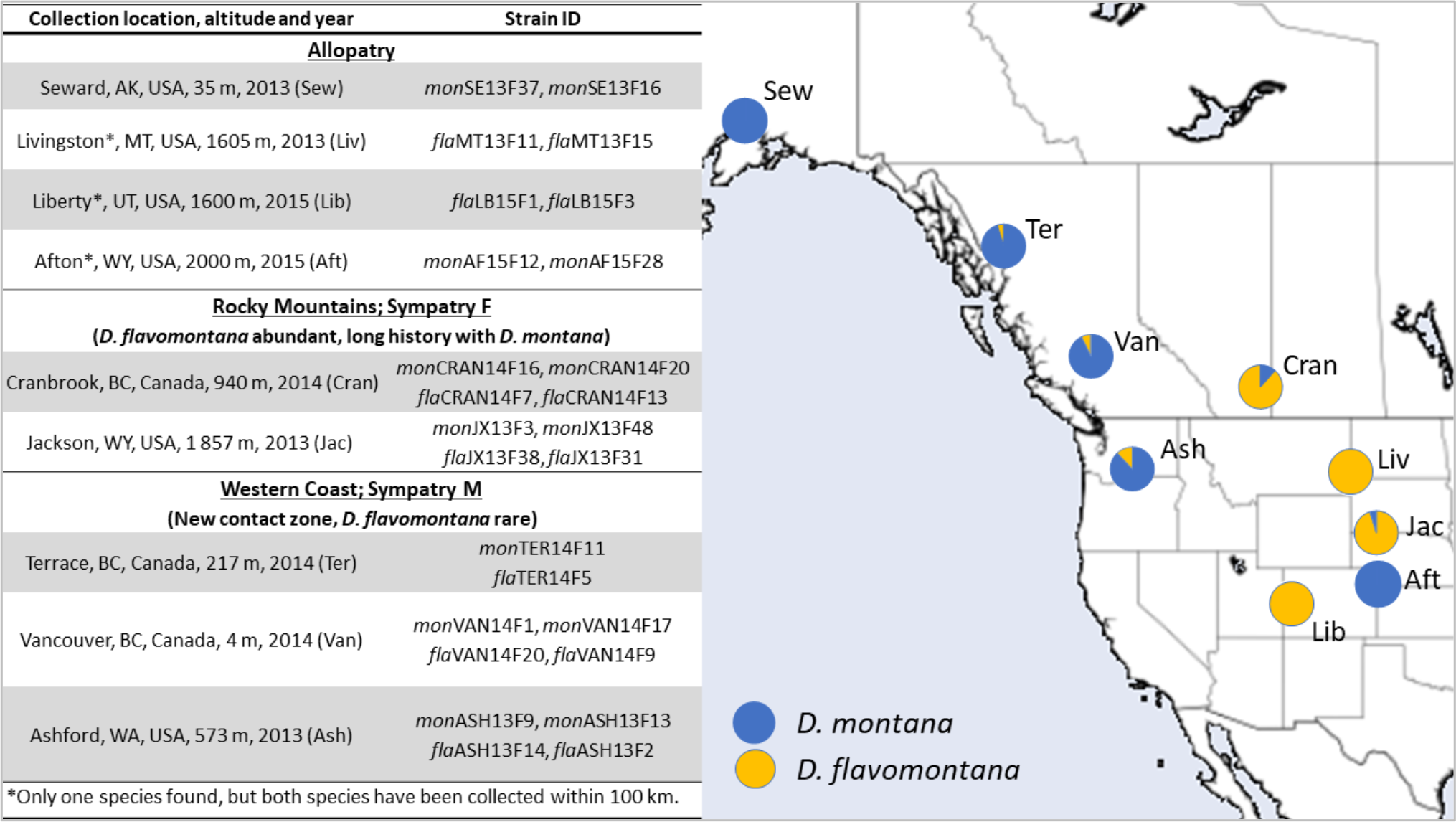
Collection sites and proportion of *D. montana* and *D. flavomontana* in North America. Sample sizes varied between ~40 and 100 individuals per site, except in Liberty where only six flies were collected. Patterson (1952) collected 203 *D. flavomontana* and only 1 *D. montana* from a large area located between Liberty and the mountain slopes inhabited by both species (Morgan district), which confirms that the population in Liberty can be regarded as an allopatric *D. flavomontana* population. Studies on reproductive barriers involved 4 strain pairs from allopatric populations, 4 pairs from Sympatry F and 5 pairs from Sympatry M (detailed information on strain pairs is given in Table S1).

#### Isofemale line establishment, maintenance and experimental use

Study strains consisted of the progenies of single overwintered, fertilized females collected in the wild in 2013 − 2015 in North America (Fig. 2, Table S1). The species of the strains was identified by sequencing part of the *COI* region in the mtDNA of one individual per progeny (see primer information in Table S2) following the protocol in Simon et al. (1994). As reproductive isolation between some Drosophila species has been found to be enhanced by Wolbachia infection (Clark et al. 2006), we tested for the presence of Wolbachia via PCR on two females and males per study strain (see detailed information in Table S2-S3) and by investigating the whole-genome sequences of four *D. montana* and five *D. flavomontana* strains. We found no evidence of Wolbachia genomic products in our samples. Therefore, any reproductive incompatibilities in our study are not explained by this endosymbiont and we do not discuss this further.

Strains were maintained and experiments performed in continuous light at 19 ± 1°C and 60-70% humidity to prevent variation in flies’ circadian rhythm and/or diapause susceptibility to affect the results. For all experiments and phenotypic assays, individuals were collected and sexed under light CO_2_ anesthesia within three days after their emergence and maintained in plastic vials containing malt-yeast medium (15-20 virgin females or males per vial). Cuticular hydrocarbons (CHCs) were extracted at the age of 14 days when the females’ ovaries can be expected to be fully developed (Salminen and Hoikkala 2013). Reproductive isolation experiments and phenotypic assays were conducted at the age of 18-22 days, when studies on *D. montana* mate choice are usually done (e.g. Jennings et al. 2014). All reproductive barriers were investigated using all strain pairs unless mentioned otherwise (Table S1).

### POSTZYGOTIC BARRIERS

Postzygotic barriers between *D. montana* and *D. flavomontana* were studied by quantifying the viability, sex ratio and fertility of hybrid offspring from reciprocal interspecific crosses. F_1_ hybrids were obtained by putting 10 females of one and 10 males of the other species into a malt vial (20 replicates for each reciprocal cross) and transferring them into a fresh vial once a week for about one month. Viability of F_1_ hybrids was determined by counting the number of 3^rd^ instar larvae and females and males that were viable at least 24 hours after emergence (note that numbers of earlier stage larvae could not be counted reliably). In intraspecific controls, 5 conspecific females and males were put into a malt vial and transferred into a new vial every day for a week to prevent overcrowding (one replicate for two strain pairs per population type), and the same traits were measured.

Interspecific F_1_ hybrids were collected from the vials within three days after eclosion. Their fertility was measured as the ability to produce progeny (at least one larva) when backcrossed to either *D. montana* or *D. flavomontana*. Each female (or male) hybrid was given a choice between males (or females) of both parental species. Hybrids that did not mate in the first trial were used in up to two more trials. Fertility of intraspecific F_1_ hybrids was studied by performing single-pair matings between F_1_ females and F_1_ males from the same cross.

All statistical analyses were conducted in R (Version 3.4.3; R Core Team 2017) and R studio (Version 1.1.383). We tested whether viability of intra- and inter-specific F_1_ hybrids varied among crosses or among population types within a cross using generalized linear mixed model (GLMM), with viability as response variable and cross or population type as an explanatory variable. These analyses were done using *glmer* function of nlme package (Pinheiro et al. 2018) and specified a binomial distribution with logit link. Strains were treated as a random effect (nested within population type and cross). In one mon♀×fla♂ cross variation of a response variable was low (excess of zeroes), and a chi squared likelihood ratio test instead of a z-test was used to test the significance. We also used one-sample student’s t tests using *t test* function of the stats package to test whether the proportion of female F_1_ hybrids differed from the expected 0.50 among crosses and population types, and whether fertility of F_1_ hybrid females and males deviated from the expected 1. Detailed statistics (degrees of freedoms, test statistics, P-values) and additional information on results are reported in Supporting Materials.

### PREMATING SEXUAL ISOLATION AND IMPORTANCE OF COURTSHIP CUES

#### Multiple-choice and no-choice tests

The magnitude of sexual isolation between *D. montana* and *D. flavomontana* was quantified using both multiple- and no-choice tests between 9 am – 11 am for each trial. For multiple-choice tests, 30 of each sex of both species were introduced into a 6cm^3^ Plexiglas mating chamber without anesthesia (see Jennings et al. 2014). Mating pairs subsequently were removed by aspiration through holes in the mating chamber walls and identified by color (*D. montana* is darker than *D. flavomontana*). In Terrace population, where the color differences were smaller, different strains were marked by feeding individuals either red- or blue-colored food, altering the colors between trials (see Wu et al. 1995). No-choice tests involved reciprocal trials of 30 females of one and 30 males of the other species. The protocol was the same as in the multiple-choice tests, except that individuals were observed for 2 hours. Controls for the no-choice tests were obtained by performing reciprocal crosses between two conspecific strains per population type (Table S1). Both multiple- and no-choice experiments were replicated five times (controls for no-choice tests one replicate), and mated females from no-choice tests were saved for PMPZ studies (see below). Sexual isolation was also studied between conspecific *D. flavomontana* strains comparing each population type using multiple-choice tests, and replicated three times, as described for heterospecific crosses (Table S1). In these experiments the flies of each strain were always marked with a different color, as explained above.

The strength of sexual isolation was calculated based on the first 50% of matings in multiple-choice tests, using the JMating 1.0.8 program (Rolan-Alvarez and Caballero 2000; Carvajal-Rodriguez and Rolan-Alvarez 2006). Here the index of sexual isolation, *I_PSI_*, is calculated from the total number of each type of mating pair, and the asymmetry index, *I_APSI_(ab/ba)*, calculates potential differences in female preference for heterotypic males in reciprocal crosses. *I_PSI_* ranges from −1 to 1, −1 denoting disassortative mating, 0 random mating and 1 complete sexual isolation, and I_APSI_ is calculated by dividing heterotypic I_PSI_-values with each other. Significance of each index was determined by bootstrapping 10 000 times in JMating.

Interspecific no-choice tests were analyzed as a proportion of mated females using generalized linear mixed model (GLMM) with binomial distribution using crosses and population types within a cross as an explanatory variable as described in the section “Postzygotic isolation” above.

#### Species differences in the importance of potential sexual cues

Contribution of visual, auditory (courtship song) and olfactory (cuticular hydrocarbons) cues in mate choice and species recognition of *D. montana* and *D. flavomontana* was determined by performing four sets of experiments with partially sensory deprived individuals within and between the species. Flies’ mating success was measured in the following treatments: (1) control - both females and males were unmanipulated and the experiment was done in light, (2) visual - both females and males were unmanipulated, but the experiments were run in darkness, (3) auditory – females were unmanipulated but males were muted by micro-scissoring off their wings, and (4) olfactory and auditory - the entire antennae of females, the third segment and aristae of which receives olfactory and auditory cues (Carlson 1996; Tauber and Eberl 2003), were removed with tweezers.

Experiments were done for one strain pair of the two species from each population type, and different experiments involving the females of the same strain were run on the same day. In each treatment and experiment, 15 females and 15 males (either conspecific or heterospecific) were placed in a vial containing malt-yeast medium. After 24 hours the females were CO_2-_anesthetized with their reproductive tracts dissected on a microscope slide in a drop of PBS-solution, covered with a cover slip, and examined under light microscopy to determine the presence of sperm.

Differences between treatments in the proportion of mated females was analyzed with generalized linear mixed model (GLMM) with binomial distribution (other details described in the “Postzygotic isolation” section above).

#### Male courtship song analysis

The songs of *D. montana* and *D. flavomontana*, produced by male wing vibration, differ clearly from each other (Päällysaho et al. 2003) and courtship songs are important in female mate choice and species discrimination (Ritchie et al. 1998; Saarikettu et al. 2005). Variation within and between the species in these cues was investigated by analyzing the songs of five males of each study strain. For song recording, a sexually mature virgin female and male of the same strain were transferred into a small petri dish, which had a moistened filter paper on the bottom and a nylon net roof. Courting males walked upside down on the roof of the chamber, which allowed song recording by holding the microphone (JVC) directly above the male. Songs were recorded using a digital Handy Recorder H4n at a temperature of 20 ± 1°C and analyzed with the Signal 4.0 sound analysis system (Engineering Design, Belmont, MA, USA). Song traits analyzed from oscillograms included the number of pulses in a pulse train (PN), the length of a pulse train (PTL), the length of a sound pulse (PL), the interpulse interval (IPI; the length of the time from the beginning of one pulse to the beginning of the next one) and the number of cycles in a sound pulse (CN; see Fig. A1). PN and PTL were analyzed for three whole pulse trains and PL, IPI and CN for the third or fourth pulse of each of these trains in oscillograms (see Fig. A1). In addition, song’s carrier frequency (FRE) was measured from the frequency spectrum of the pulse trains.

Mean values of song traits were averaged over three pulse trains of each male. To reduce the number of variables in the dataset, a principal component analysis (PCA) was applied using the *prcomp* function in R (Version 3.4.3) and R studio (Version 1.1.383). Before running the PCA, pulse train length (PTL) was removed from the analysis due to its strong correlation with pulse number (PN) both in *D. montana* (84%) and *D. flavomontana* (85%). PCA scores for each study strain were centered and scaled. The differences in each courtship parameter within species between population types were analyzed with linear mixed model (LMM) using study strains as a random effect. These analyses were done using *lmer* function of nlme package (Pinheiro et al. 2018).

#### Cuticular hydrocarbon (CHC) profiles

CHCs may serve as contact pheromones and function in female discrimination of mates (Ferveur 2005; Jennings et al. 2014a). CHC profiles were analyzed for both sexes of all study strains (usually 5 individuals/sex/strain; Table S4). CHC extractions were performed in the morning by immersing individuals in 200 µl of n-hexane in glass vials (Micro Liter Analytical Supplies; 1.8 ml) for 10 min, after which individuals were removed. Open vials were maintained in a sterile fume hood at room temperature until the hexane had evaporated, then vials were sealed and stored at −20°C. Control vials with pure solvent (n-hexane) were prepared in the same way.

CHC extracts were analysed with an Agilent7890 gas chromatograph (GC) coupled with an Agilent 5975C Mass Selective (MS) Detector (Agilent, Waldbronn, Germany) at the University of Wuerzburg (Germany). The GC (split/splitless injector in splitless mode for 1 min, injected volume: 1 µl at 300°C) was equipped with a DB-5 Fused Silica capillary column (30m x 0.25 mm ID, df = 0.25 µm; J&W Scientific, Folsom, USA). Helium served as a carrier gas at a constant flow of 1 ml/min. The temperature program consisted of the start temperature 60°C, temperature increase by 5°C/min up to 300°C and maintenance at 300°C for 10 min. The electron ionization mass spectra (EI-MS) were acquired at an ionization voltage of 70 eV (source temperature: 230°C). Chromatograms and mass spectra were recorded and quantified with the software Agilent Enhanced Chem Station G1701AA (version A.03.00). Individual CHC compounds were chemically identified using the MS data base Wiley275 (John Wiley & Sons, New York, USA), retention indices, and the detected diagnostic ions (Bernier et al. 1998). Some substances could not be accurately separated and, in these cases, the combined quantity was calculated by integrating over all substances within a peak.

CHC profile similarity was assessed by means of multivariate Linear Discriminant Analysis (LDA) and Random forest classification using the functions *lda* of MASS package and *randomForest* of randomForest package (Liaw and Wiener 2002). In addition, Bray-Curtis dissimilarities were analyzed for differences between species in each population type and differences between sexes within a population type for both species. Values range from 0 to 1, where 0 means the same composition and 1 means complete dissimilarity. Significance levels were tested with linear mixed model (LMM) using study strains as a random effect.

### POSTMATING-PREZYGOTIC (PMPZ) BARRIERS

Females that copulated for at least 3 minutes (to ensure sperm transfer: Mazzi et al. 2009) with a heterospecific male were obtained from the no-choice tests (described above, section “Multiple-choice and no-choice tests”). As the number of matings between *D. montana* females and *D. flavomontana* males was low, we generated more matings in this direction by playing females conspecific song (see e.g. Saarikettu et al. 2005) while being exposed to *D. flavomontana* males that were muted (described above, section “Species differences in the importance of potential sexual cues”) 1d before the mating experiment. *D. montana* females and muted *D. flavomontana* males (n=10-15 per trial) were placed in a mating arena (small petri dish and a nylon net roof) placed above a subwoofer (Harman Kardon JBL Platinum Series Speakers) connected to a computer. Recorded *D. montana* song was played throughout the courtship, and mating pairs were collected once copulation had ended. 10 reciprocal single-pair crosses were made between the females and males of two conspecific strains from each location for intraspecific controls.

PMPZ barriers were quantified by assessing sperm transfer and storage, and the production and fertilization of eggs in all interspecific crosses and their controls. Mated females were placed individually into a set of 20 vials (“manifold”) with 1 cm of malt-yeast medium at the bottom and dissected after 48 hours to check for the presence of sperm in their seminal receptacles and spermathecae. The amount of sperm was estimated using four categories: 0 = no motile sperm, 1 = maximum of two sperm cells, 2 = intermediate amount of sperm and 3 = seminal receptacles and/or spermatheca full of sperm. The number of eggs laid by each female in the vial was counted immediately after her removal, and again after three days, to calculate the proportion of eggs that had hatched and proceeded to larval stage during this period. Finally, as also virgin females lay eggs, we asked whether and how much the receipt of sperm increases females’ fecundity.

Reduction in the proportion of hatched eggs may result from either fertilization failure (PMPZ barriers) or from problems in embryo development due to genetic incompatibilities (postzygotic barriers). To distinguish between these, egg fertilization and embryo development were investigated in eggs laid by *D. flavomontana* females that had mated to *D. montana* males (reciprocal cross was not studied because *D. montana* females did not store *D. flavomontana* sperm), and between *D. flavomontana* populations, using all strain pairs. Freshly laid eggs of 17-33 mated females per strain were collected each day for 3d, then fixed and processed for fluorescence microscopy (DAPI, Snook and Karr 1998; Jennings et al. 2014b). Eggs were classified as developing if either clear mitotic division or cellular differentiation was evident. Eggs that did not meet these criteria (Fig. A2) were examined for the presence of sperm inside the egg to determine whether these were fertilized but karyogamy had not yet occurred or whether they were unfertilized (i.e. sperm were absent). The presence of sperm inside eggs was scored using differential interference contrast (DIC) light microscopy (Jennings et al. 2014b). Sperm length of *D. montana* is 3.34 ± 0.02 mm and of *D. flavomontana* is 5.53 ± 0.01 mm (Pitnick et al. 1999), thus the sperm flagellum can easily be seen as a coiled structure near the anterior end of the egg (see Fig. A2).

Variation in females’ sperm storage ability among intra- and inter-specific crosses and between population types within a cross was tested treating this trait as an ordinal variable in cumulative link mixed model (CLMM). These analyses were conducted using *clmm* function of ordinal package (Christensen 2018). Proportion of hatched/developing eggs were analyzed as in the “Postzygotic isolation” section, using generalized linear mixed model (GLMM) with binomial distribution. In some mon♀×fla♂ crosses, variation of a response variable was low (excess of zeroes), and a chi squared likelihood ratio test instead of a z-test was used to test the significance. Finally, we tested whether the presence of sperm had increased number of eggs laid (fecundity) in each cross using generalized linear mixed model (GLMM) with negative binomial distribution, with fecundity as a response variable, presence of sperm (yes/no) as an explanatory variable and study strains as random effects. These analyses were carried out using *glmmadmb* function of glmmADMB package (Skaug et al. 2013).

## Results

### POSTZYGOTIC BARRIERS – FITNESS OF F_1_ HYBRIDS

We first determined the strength of postzygotic isolation to define the cost of interspecific matings, which generates selection for reinforcement, by producing intra- and inter-specific F_1_ hybrids and measured their viability (from the 3^rd^ instar larvae to adult), sex ratio, and fertility. We found that crosses between *D. flavomontana* females and *D. montana* males produced a higher number of 3^rd^ instar larvae than the reciprocal cross, especially in Allopatry (339 and 31 larvae, respectively; Fig. 3A-B), which could be due to that the flies had not mated or that the females had problems in sperm usage. Viability of F_1_ hybrids from crosses between *D. montana* females and *D. flavomontana* males was very low compared to that of the intraspecific crosses, except in Sympatry F (Fig. 3C). Due to small sample sizes, sex-ratio bias could not be tested statistically in these interspecific crosses. Viability of F_1_ hybrids from crosses between Allopatric *D. flavomontana* females and *D. montana* males did not differ from viability of intraspecific progeny, while in Sympatry F or Sympatry M F_1_hybrid viability was significantly lower than in intraspecific crosses (Fig. 3D, Table S5). The opposite pattern was seen for sex-ratio bias: compared to intraspecific crosses, F_1_ hybrids between Allopatric *D. flavomontana* females and *D. montana* males were significantly female-biased whereas there was no significant difference in sex ratio of F_1_ hybrids arising from either Sympatry F or Sympatry M (Fig. 3B, Table S5).

**Figure 3.**
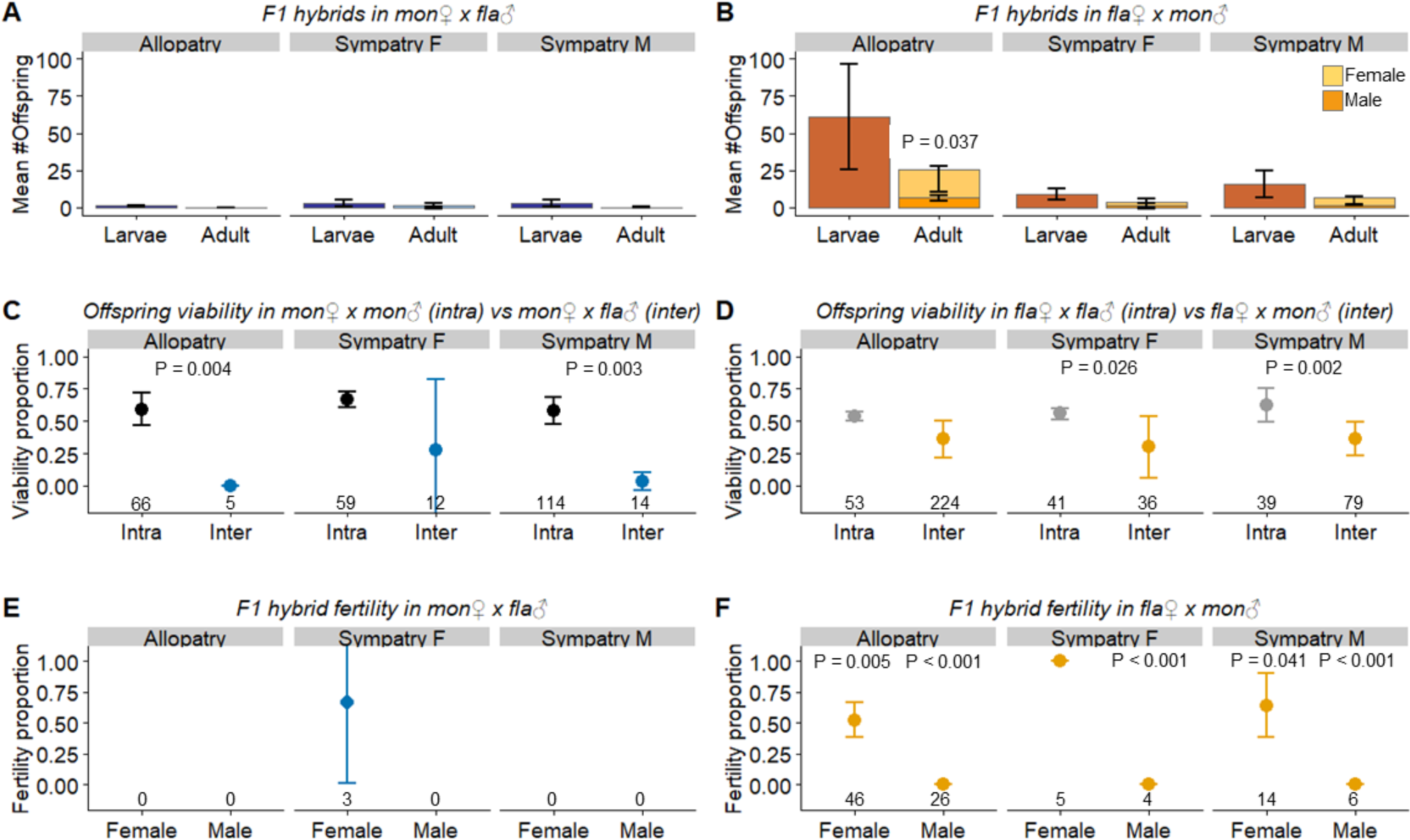
Offspring production and viability in intra- and inter- specific crosses and fertility of F_1_ hybrids in different population types. Error bars represent 95% confidence intervals. (A) F_1_ hybrid larvae and adults produced by *D. montana* females and *D. flavomontana* males. (B) F_1_ hybrid larvae and adults (females and males) produced by *D. flavomontana* females and *D. montana* males. (C) Viability of F_1_ hybrids produced in intra- and inter-specific crosses involving *D. montana* females. (D) Viability of F_1_ hybrids in intra- and inter-specific crosses involving *D. flavomontana* females. Differences between intra- and inter-specific crosses were significant in both sympatric populations. (E) Fertility of F_1_ hybrids produced in interspecific crosses involving *D. montana* females (could not be statistically tested due to small sample size). (F) Fertility of F_1_ hybrids produced in interspecific crosses involving *D. flavomontana* females (could not be statistically tested for Sympatry F females since all of them were fertile). The numbers above x-axis refer to the total number of studied larvae (C and D) or adult flies (E and F).

To determine F_1_ hybrid fertility, hybrids were backcrossed to either of the parental species. There was no effect of parental species on F_1_ hybrid fertility (GLMM, z_1,99_ = 1.21, P = 0.228) so subsequent statistics were performed on combined data of these crosses. Among the few hybrids produced in crosses between *D. montana* females and *D. flavomontana* males, only three females mated and two of them were fertile (Fig. 3E). Crosses between *D. flavomontana* females and *D. montana* males produced 106 F_1_ hybrids and, while 101 mated to parental species (Table 1), all F_1_ males were sterile and at least half the F_1_ females fertile (Fig. 3F).

**Table 1.** The most influential CHC substances based on random forest analysis (see Fig. S2). Most of the compounds included alkenes with different numbers of carbons and different double-pond positions in a chain, while in *D. flavomontana* sympatric populations class of 2-methyl-branched alkanes and/or alkadienes have a large contribution on sex differences.

| Random forest analysis | Allopatry | Sympatry F | Sympatry M |
| --- | --- | --- | --- |
| Between *D. montana* and *D. flavomontana* | 2methylC24-alkane<br>C27-alkene 3 | C27-alkene 1<br>C27-alkene 3 | C27-alkene 2<br>2methylC24-alkane |
| Between sexes of *D. montana* | C25-alkene 2<br>C29-alkene 1 | C25-alkene 4<br>C29-alkadiene 2 | C27-alkene 2<br>C29-alkene 2 |
| Between sexes of *D. flavomontana* | C27-alkene 5<br>C25-alkene 4 | 2methylC28-alkane/C29-alkadiene 5<br>C27-alkene 2 | 2methylC28-alkane/C29-alkadiene 5<br>2methylC30-alkane/C31-alkadiene 4 |

### SEXUAL ISOLATION AND THE FACTORS MAINTAINING IT

#### The strength and asymmetry of sexual isolation between the species and between D. flavomontana populations

The strength of sexual isolation between *D. montana* and *D. flavomontana* was studied using multiple-choice and no-choice tests, and between *D. flavomontana* populations with multiple-choice tests. In interspecific multiple-choice tests, matings occurred mainly within the species and the sexual isolation index (I_PSI_) varied from 0.89 to 0.92 (Fig. 4A, Table S6). The asymmetry index (I_APSI_) showed that *D. flavomontana* females and *D. montana* males mated more than the flies of the reciprocal cross when individuals were from either Allopatry or Sympatry M, but not Sympatry F (Fig. 4B, Table S6; see data for individual strain pairs in Table S7).

**Figure 4.**
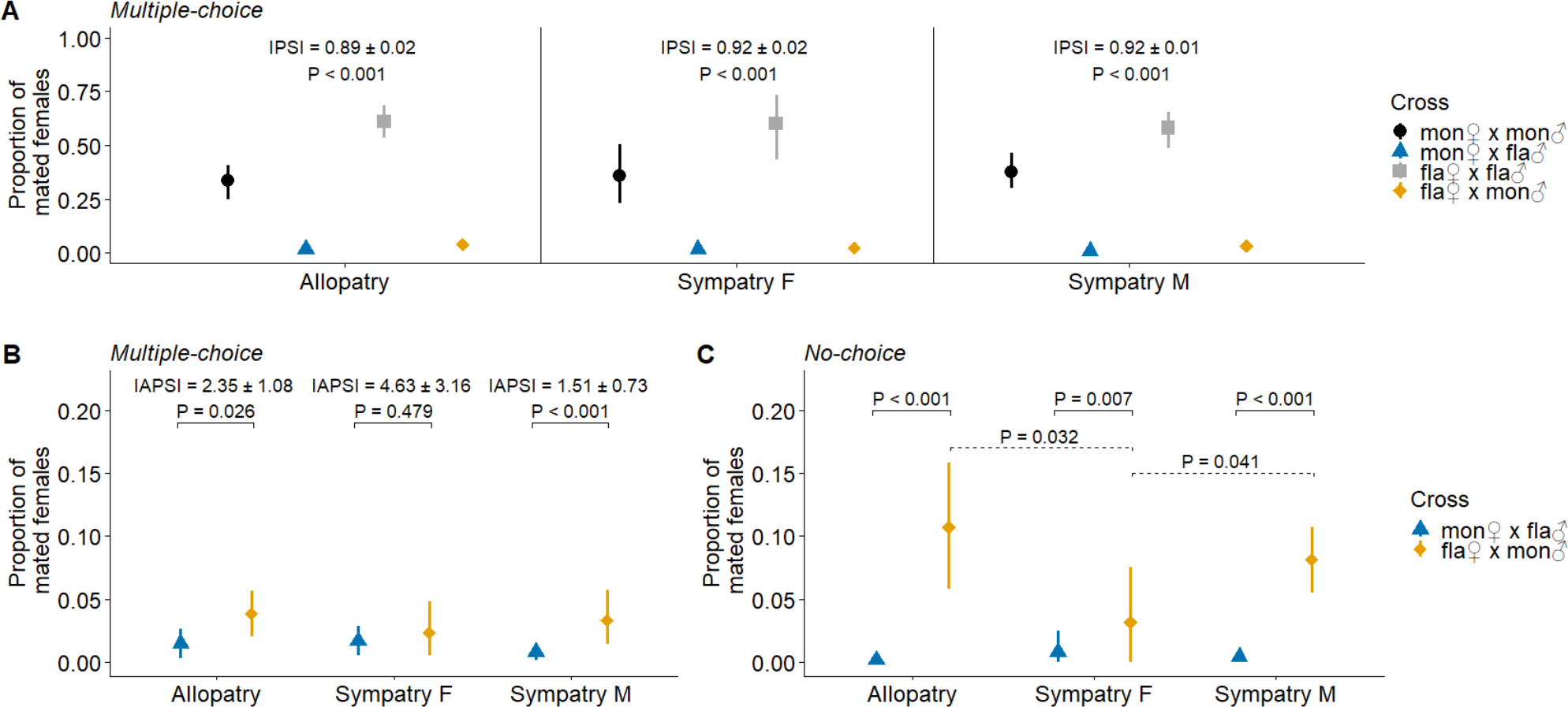
The strength of sexual isolation between *D. montana* and *D. flavomontana* in multiple- and no-choice tests for different population types. (A) Sexual isolation index (I_PSI_) was significant, and biased towards intraspecific matings, in all population types. (B) Multiple-choice asymmetry index (I_APSI_) tests showed that population type impacted asymmetry: *D. flavomontana* females and *D. montana* males mated more often than the reciprocal when individuals arose in Allopatry and Sympatry M, but not in Sympatry F. (C) No-choice tests showed that *D. flavomontana* females and *D. montana* males mated more than the flies of the reciprocal in all population types (solid line P values). Also, *D. flavomontana* females were more likely to mate with heterospecific males in Allopatry and Sympatry M than in Sympatry F (dashed line P values). Error bars represent bootstrapped 95% confidence intervals.

No-choice tests measured the strength of sexual isolation when individuals were offered only heterospecific mating partners. In all population types, the proportion of mated *D. flavomontana* females was higher than that of *D. montana* females (Fig. 4C, Table S6), indicating that *D. flavomontana* females are less discriminating than *D. montana* females against heterospecific males. The proportion of mated *D. montana* females was equally low in all population types (Table S6), while *D. flavomontana* females from Allopatry and Sympatry M mated more frequently with heterospecific males than the ones from Sympatry F (Fig. 4C, Table S6). However, the proportion of mated females remained very low in both interspecific crosses (0.00-0.01 between *D. montana* females and *D. flavomontana* males and 0.03-0.11 between *D. flavomontana* females and *D. montana* males; see Fig. 4C) compared to intraspecific crosses (*D. montana*: 0.90-0.97; *D. flavomontana:* 0.82-0.92).

The strength of sexual isolation between *D. flavomontana* populations, measured in multiple-choice tests, revealed significant isolation between individuals from the Sympatry M population type crossed with individuals from the Sympatry F population type (Fig. 5; see data for individual strain pairs in Table S8). This isolation was asymmetrical, with females from Sympatry F mating less often with males from Sympatry M than vice versa. Other comparisons showed no sexual isolation.

**Figure 5.**
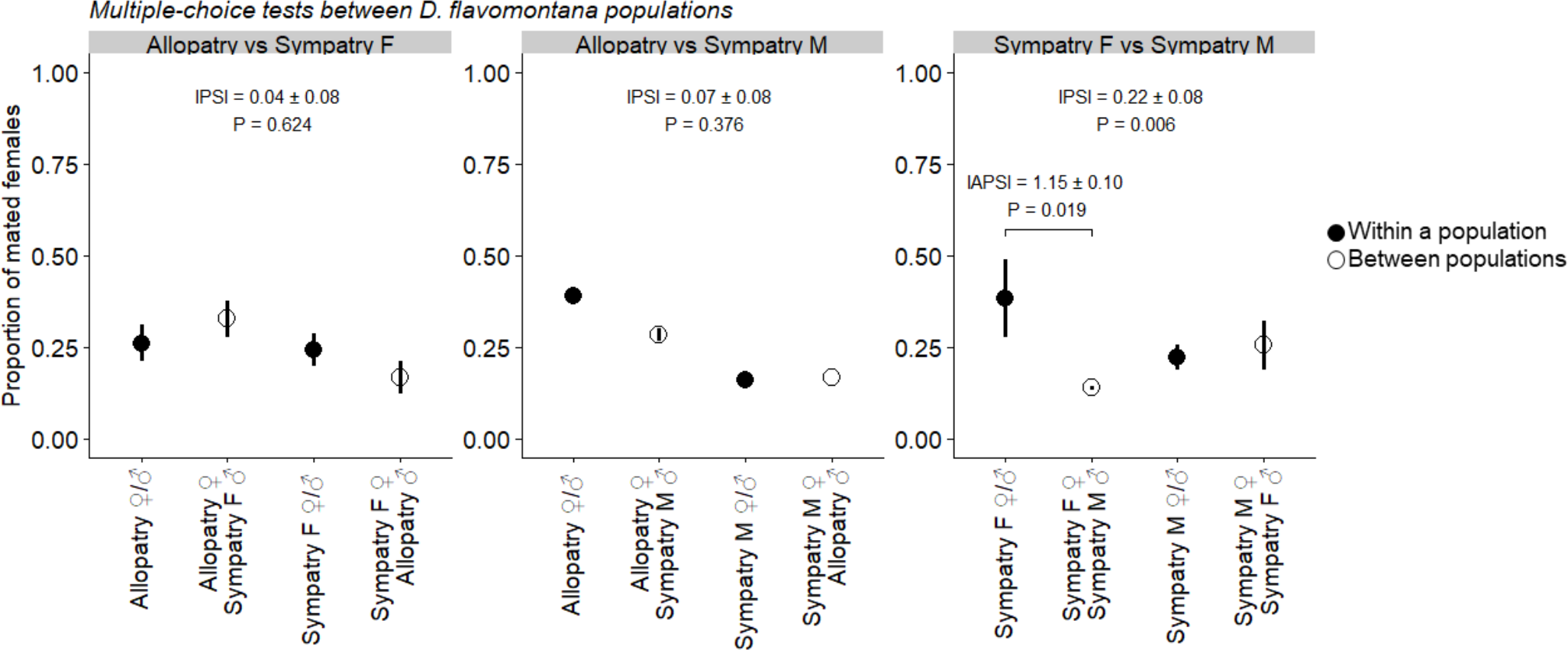
The strength of sexual isolation between different *D. flavomontana* populations (multiple-choice test). Females from Sympatry F showed significant discrimination against males from Sympatry M, but the other comparisons showed no sexual isolation. Error bars represent bootstrapped 95% confidence intervals.

#### Importance of sexual cues in species recognition / sexual selection

The importance of visual, auditory and olfactory cues in species-recognition and/or sexual selection was studied by comparing flies’ mating propensity between the control trial and the test trials where the transmission of one or more cues was prevented. Visual signals did not play an essential role in mating success in either species, as mating frequency was at the same level in light (control) and in dark (Fig. 5A and C, Table S9). However, the two species differed in the impact of auditory and olfactory signals on mating success. *D. montana* females did not mate without hearing species-specific male courtship song (Fig. 6A, Table S9) whereas *D. flavomontana* females mated equally often with control and wingless (muted) males of their own species (Fig. 6C, Table S9). Removal of female antennae, which silenced both auditory and volatile olfactory cues, prevented mating in *D. montana*, as expected given the results of auditory manipulation alone (Fig. 6A, Table S9), and significantly reduced male *D. flavomontana* mating success (Fig. 6C, Table S9).

**Figure 6.**
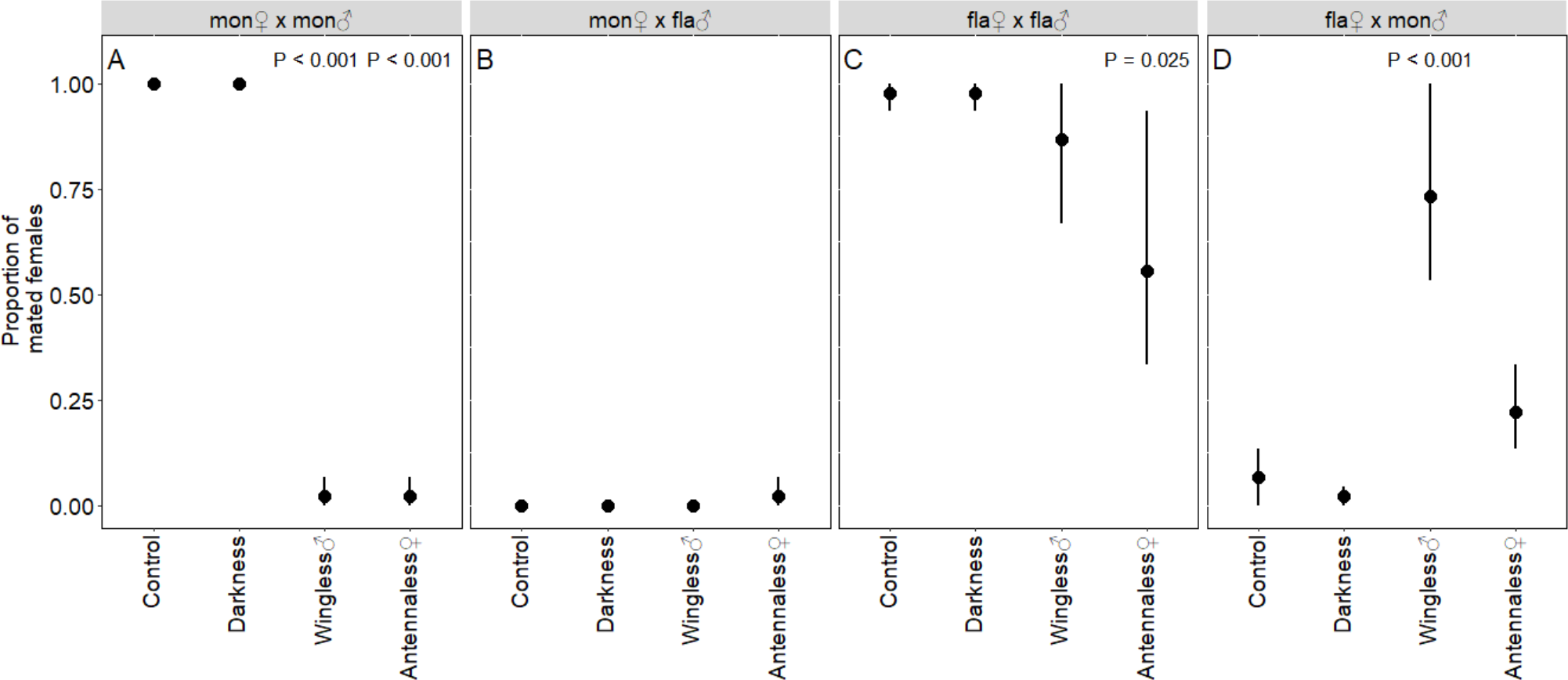
The impact of blocking the transfer of sensory cues on the proportion of mated females in intra- and inter-specific crosses. (A) *D. montana* females did not mate with conspecific males when the wings of the males or the antennae of the females had been removed. (B) *D. montana* females did not mate with *D. flavomontana* males, except once when female antennae were removed. (C) *D. flavomontana* females mated significantly less with conspecific males when female antennae were removed. (D) *D. flavomontana* females mated significantly more with wingless (muted) *D. montana* males than with unmanipulated (control) males. Error bars represent bootstrapped 95% confidence intervals.

Nearly all interspecific matings occurred between *D. flavomontana* females and *D. montana* males in trials where male wings or female antennae had been removed (Fig. 6D, Table S9). *D. flavomontana* females mated significantly more with wingless than with normal *D. montana* males (Fig. 6D, Table S9), i.e. hearing a heterospecific song decreased their willingness to mate more than silence. Overall, these results suggest that *D. montana* require male song (and perhaps CHCs) whereas the courtship of *D. flavomontana* relies more on CHCs.

#### Divergence in important sexual cues within and between species

Divergence of important sexual cues, including male courtship song and CHCs, was studied between conspecific populations and between species. Variation in male song traits within and between species is illustrated with a principal component analysis plot (Fig. 7B). The first two components accounted for 84.5% of the total between-male variance (Fig. S1, Table S10). The first principal component separated pulse number (PN), pulse length (PL) and number of cycles per pulse (CN) from interpulse interval (IPI) and explained 61.0% of the variation (Fig. 7B). The second principal component explained 23.5% of variation, and here the number of cycles per pulse (CN) varied both within and between species, while the song frequency (FRE) varied only within *D. montana*. In *D. montana* CN and FRE were slightly higher in males from Allopatry than in males from Sympatry M (LMM, CN: t_1,43_ = −3.04, P = 0.019; FRE: t_1,43_ = −2.45, P = 0.040), while none of the *D. flavomontana* song parameters varied significantly between population types (Table S11-12).

**Figure 7.**
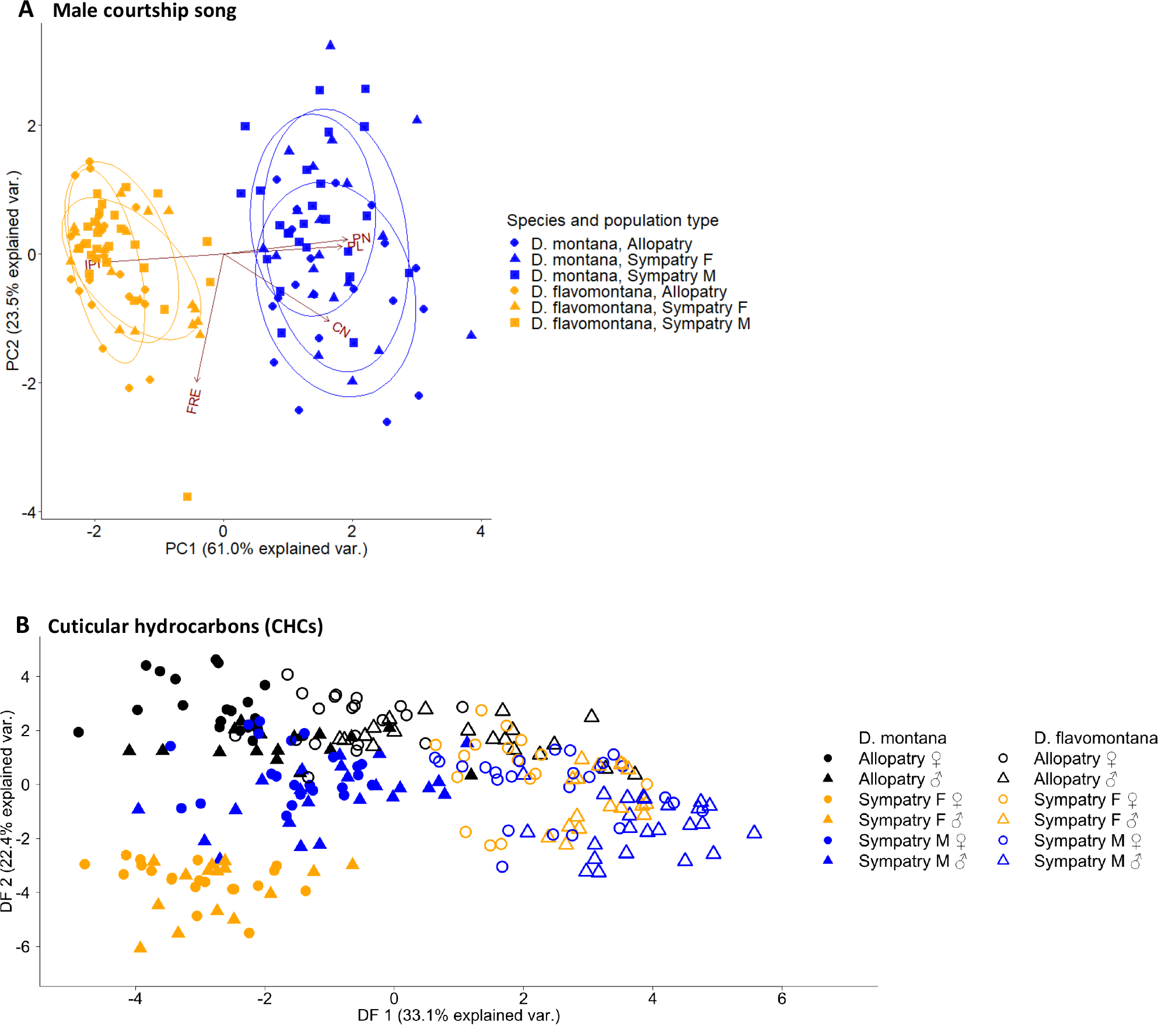
Variation between species and population types in male song and the CHCs of both sexes. (A) Male courtship songs showed clear divergence between the species, while differences within the species were relatively small. (B) CHCs varied both within and between the species with species differences being greatest in sympatric populations. In addition, sex differences were higher within *D. flavomontana* than within *D. montana*.

CHCs of allopatric *D. montana* and *D. flavomontana* populations resembled each other more than those from either of the sympatric population types (Fig. 7C). Species differences, calculated as Bray-Curtis dissimilarities, were significantly higher in Sympatry F (0.52 ± 0.11) and in Sympatry M (0.51 ± 13) than in Allopatry (0.36 ± 0.10; LMM, _t1,2670_ = 6.60, P < 0.001 and _t1,3783_ = 5.81, P < 0.001,respectively), while Sympatry F and Sympatry M did not differ significantly from each other (LMM, _t1,3491_ = −0.39, P = 0.697).

Within *D. montana*, CHC differences between sexes were higher in Sympatry M (0.39 ± 0.14) than in Allopatry (0.30 ± 0.12, LMM, _t1,910_ = 2.52, P = 0.016), but equally high with Sympatry F (0.31 ± 0.13, LMM, _t1,850_ = 1.05, P = 0.299). They did not differ between Allopatry and Sympatry F either (LMM, _t1,658_ = 1.09, P = 0.284). Within *D. flavomontana*, CHC differences between sexes were more pronounced in Sympatry M (0.51 ± 0.13) than in Allopatry (0.37 ± 0.10; LMM, t_1,978_ = 4.13, P < 0.001) or in Sympatry F (0.41 ± 0.10; LMM, t_1,887_ = 2.30, P = 0.027), where they were of the same level (LMM, t_1,667_ = 1.63, P = 0.113). Overall sex differences were higher in *D. flavomontana* than in *D. montana* (LMM, _t1,2479_ = −4.55, P < 0.001), further indicating that CHCs are more important in the sexual selection and/or species-recognition of *D. flavomontana* than that of *D. montana*. Confusion matrix for Random forest analysis showed only a few classification errors beyond the population type (Table S13). The sexes were confused slightly more often in *D. montana* than in *D. flavomontana*, in congruency with the higher chemical differentiation of the sexes in the latter species.

The most influential CHC substances for the chemical dissimilarities between species and between sexes within species in each population type were defined using random forest analysis (Table 1, Fig. S2). Most of the substances were alkenes with varying numbers of carbons in a chain and with different double-pond position. Interestingly, in both sympatric *D. flavomontana* population types, 2-methyl-branched alkanes and/or alkadienes had a large contribution to sex differences, which indicates a signal function of these compound classes. The relative amounts of these compounds were higher in males than in females (Table S14).

### POSTMATING PREZYGOTIC (PMPZ) BARRIERS

The strength of PMPZ barriers was determined by quantifying the amount of sperm in female sperm storage organs after mating, as well as female fecundity and egg fertilization in intra- and inter-specific crosses. The presence and amount of sperm in female sperm storage organs depends on whether sperm is transferred during mating and/or whether females store and/or deplete sperm. All *D. montana* females had fewer sperm when mating with heterospecific than conspecific males (Fig. 8A, Table S15). Allopatric and Sympatry F *D. flavomontana* females stored sperm equally well regardless of whether it was received from con- or heterospecific males, while Sympatry M females had fewer sperm when mating with heterospecific than conspecific males (Fig. 8A, Table S15). Also, in interspecific crosses, sperm was more successfully transferred and stored in *D. flavomontana* than in *D. montana* females in all population types (Fig. 8A, Table S15).

**Figure 8.**
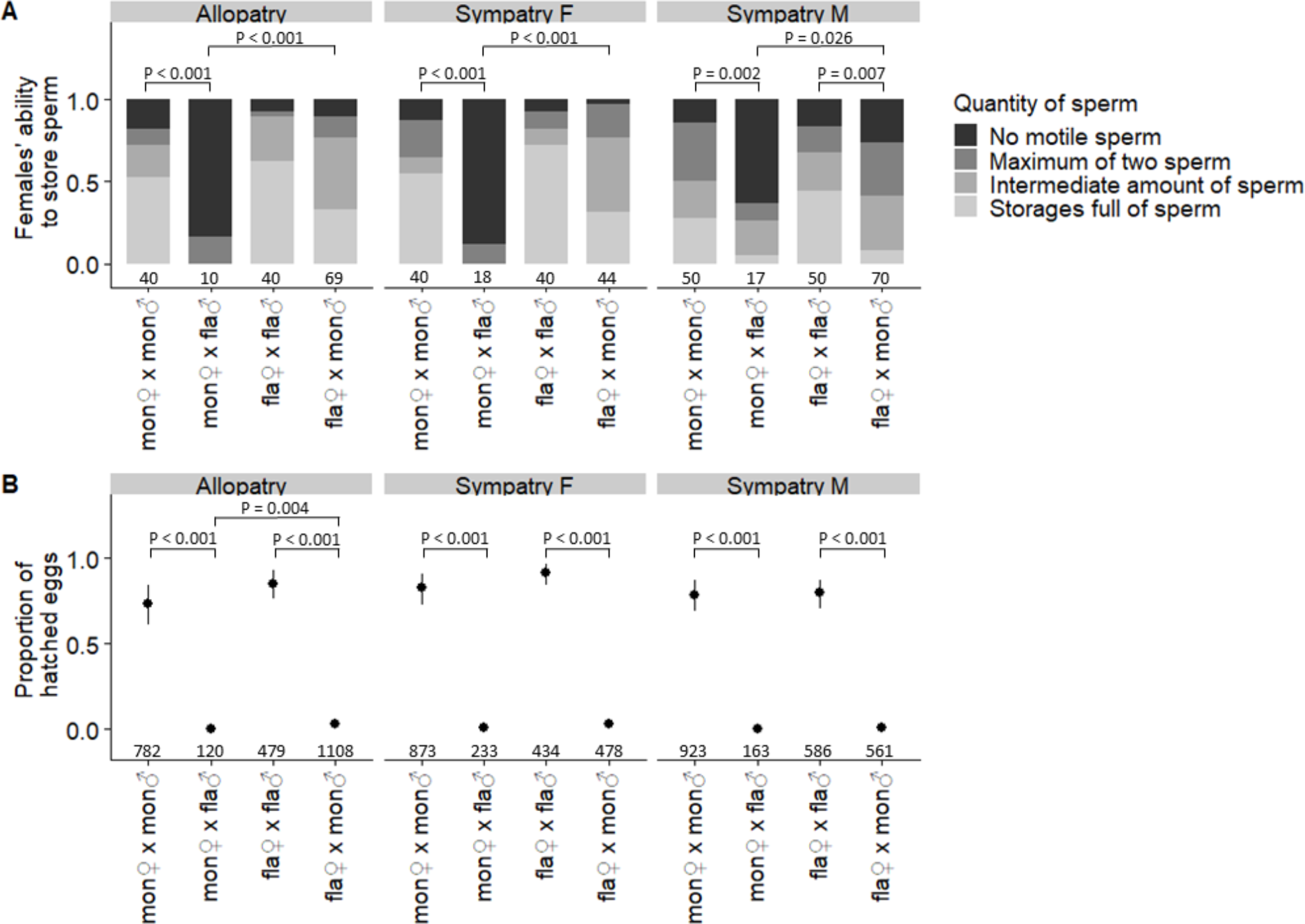
PMPZ barriers in the transfer and/or storage of sperm and the proportion of hatched eggs. (A) The transfer and/or storage of sperm was lower in all crosses between *D. montana* females and *D. flavomontana* males compared to intraspecific controls or reciprocal crosses. Sympatry M *D. flavomontana* females had fewer sperm when mating with heterospecific than conspecific males, but Allopatric and Sympatry F females stored both hetero- and con-specific sperm equally well. Numbers above x-axis refer to the number of studied females in each cross. (B) Proportion of hatched eggs was lower in all interspecific crosses than in intraspecific ones, and in Allopatry it was lower between *D. montana* females and *D. flavomontana* males than in the reciprocal ones. Numbers above x-axis refer to the number of studied eggs in each cross. Error bars represent bootstrapped 95% confidence intervals.

Sexually mature *D. montana* and *D. flavomontana* females can lay eggs as virgins. We tested whether presence of sperm (yes, no) increases female egg laying in intra- and inter-specific crosses. Statistical tests were done to the combined data without population type division since it did not explain the data statistically. Presence of sperm in intraspecific matings increased females’ average egg production in both species (*D. montana -* 13 ± 1 (sperm absent) to 21 ± 2 (sperm present) eggs, GLMM, z_129_ = 2.02,P = 0.044; *D. flavomontana* - 5 ± 1 (sperm absent) to 12 ± 2 (sperm present) eggs, GLMM, z_129_ =2.54, P = 0.010). However, in interspecific matings, male sperm did not increase fecundity in either species (GLMM, *D. montana*: z_45_ = 0.41, P = 0.680; *D. flavomontana*: GLMM, z_182_ = 1.64, P = 0.100).

The proportion of hatched eggs in reciprocal crosses between species was low (*D. montana* females and *D. flavomontana* males = 0.00-0.01; *D. flavomontana* females and *D. montana* males = 0.01-0.03) and significantly lower than in intraspecific crosses (*D. montana* = 0.73-0.83; *D. flavomontana* = 0.80-0.91; Fig. 8B, Table S15). Also, in interspecific crosses, proportion of hatched eggs was higher in *D. flavomontana* than in *D. montana* females in Allopatry, but not in sympatries (Fig. 8A, Table S15).

The low proportion of hatched eggs in crosses between *D. flavomontana* females and *D. montana* males was due to fertilization failure rather than fertilization followed by genetic incompatibility (Fig. 9). On average, only 1.3 – 5.1 % of the eggs were developing and eggs that failed the development criteria did not contain sperm. This PMPZ barrier was 4.1 and 3.9 times stronger in Sympatry M than in either Allopatry or Sympatry F, respectively, but Sympatry F did not differ from Allopatry (Fig. 9, Table S16).

**Figure 9.**
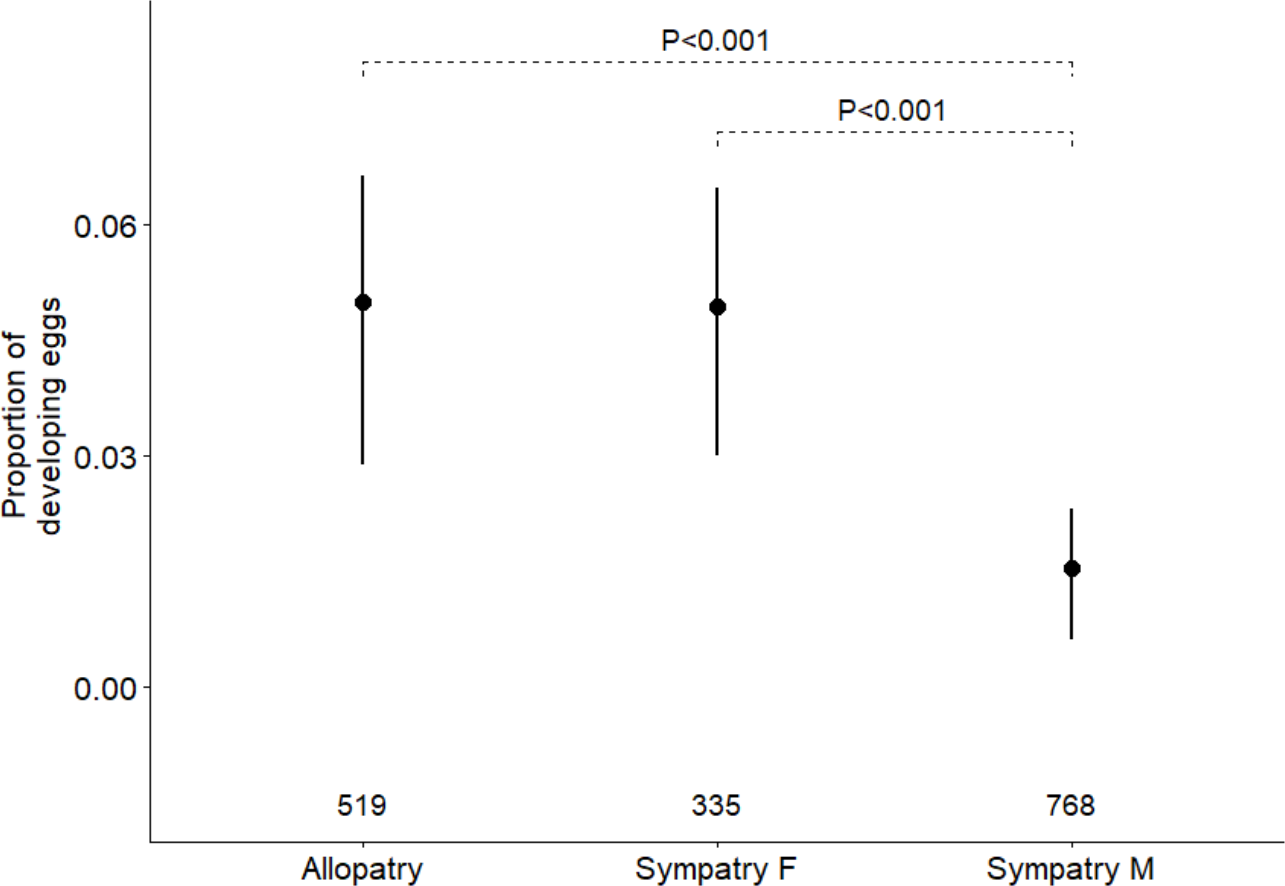
In crosses between *D. flavomontana* females and *D. montana* males, DAPI staining and microscopy revealed significantly stronger PMPZ isolation at the level of egg fertilization and development in Sympatry M compared to Allopatry or Sympatry F. Numbers above x-axis represent the number of studied eggs. Error bars represent bootstrapped 95% confidence intervals.

Reinforcement of PMPZ barriers was also detected in crosses between different types of *D. flavomontana* populations. Proportion of developing eggs was significantly reduced in crosses between Sympatry F females mated to Sympatry M males, and those between Sympatry M females and Allopatric males, compared to the other crosses for the given female types (Fig. 10, Table S17). As in interspecific crosses, none of the non-developing eggs contained sperm.

**Figure 10.**
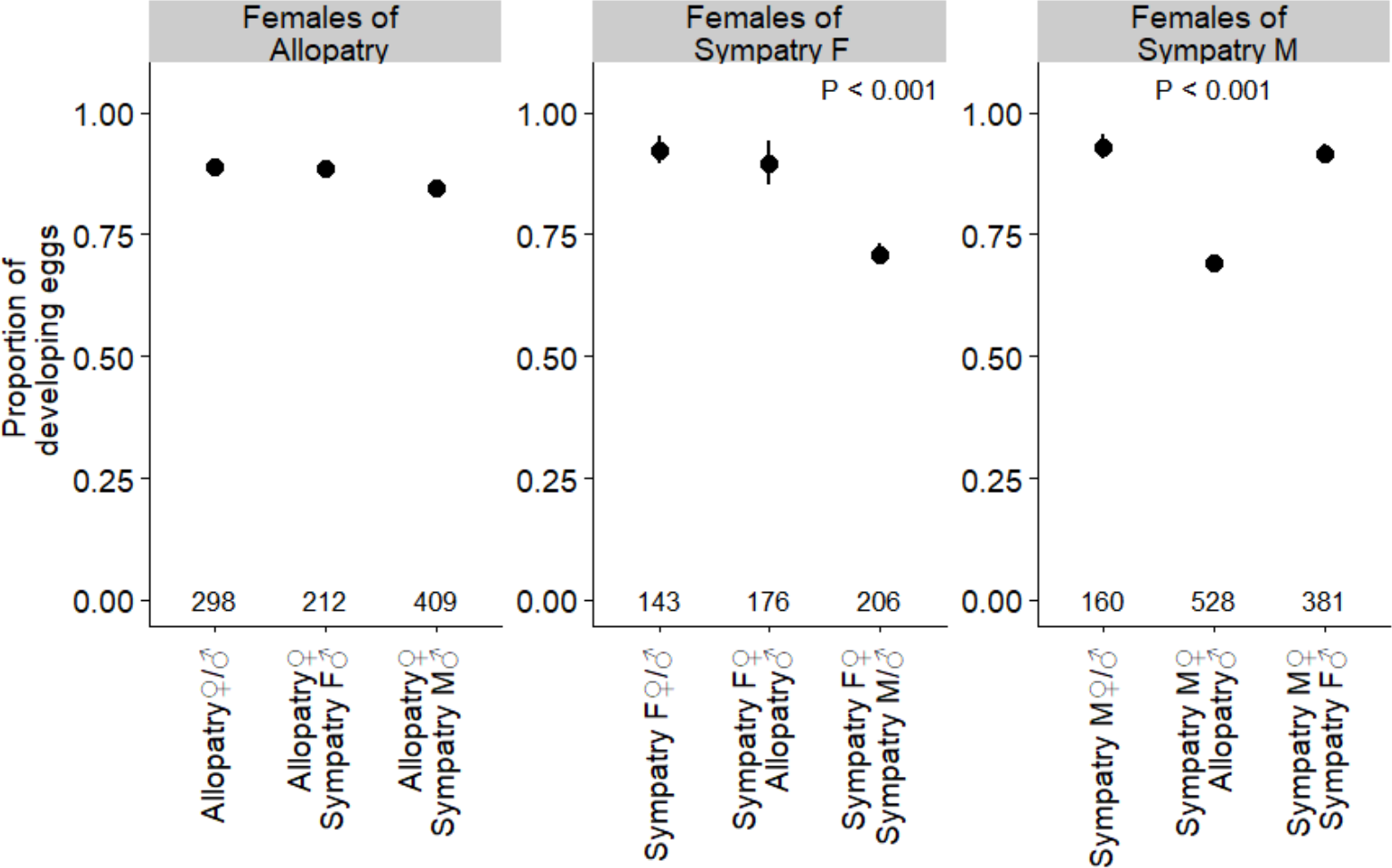
Proportion of developing eggs in crosses between *D. flavomontana* flies from different population types. Numbers above x-axis represent the number of eggs examined. Error bars represent bootstrapped 95% confidence intervals.

### THE REINFORCEMENT OF PREZYGOTIC BARRIERS

Sexual and PMPZ barriers between *D. montana* females and *D. flavomontana* males were almost complete in all population types, leaving no space for reinforcement to occur. To find out whether the costs involved in matings between *D. flavomontana* females and *D. montana* males had reinforced prezygotic reproductive barriers in sympatric populations and promoted character displacement between *D. flavomontana* populations, RI (Reproductive Isolation; Sobel and Chen 2014) was calculated separately for sexual and PMPZ isolation:

> *RI_4A_ = 1 – 2 × (H / (H + C))*
>
> *where H stands for heterospecific and C for conspecific cases*

The strength of sexual isolation between *D. flavomontana* females and *D. montana* males was calculated from no-choice results, as these reflect female discrimination and the strength of isolation better than multiple-choice tests (flies are not influenced by conspecific mating cues from other courting pairs). The strength of sexual isolation between *D. flavomontana* populations was calculated from multiple-choice results (Table S8). Calculation of the strength of PMPZ isolation was based on the proportion of developing eggs, which includes failures in sperm transfer and/or storage and egg fertilization. RI-values obtained for individual strain pairs were averaged to produce a joint value in each population type.

Sexual and PMPZ isolation between *D. flavomontana* females and *D. montana* males varied by population type (Fig. 11A). Sexual isolation was stronger in Sympatry F than in either Allopatry or Sympatry M whereas PMPZ isolation was stronger in Sympatry M than in either Allopatry or Sympatry F. Thus, in crosses where sexual isolation is less effective, PMPZ barriers could block interspecific gene flow. Reinforcement of sexual isolation had far-reaching effects in promoting sexual isolation between *D. flavomontana* populations, Sympatry F females discriminating against males of the other populations (Fig. 11B). Similarly, some populations pairs including Sympatry M individuals had slightly increased PMPZ isolation compared to other population pairs (Fig. 11B). Reduction in gene flow resulting from prezygotic isolation (composite index) was calculated as in Turissini et al. (2015):

> *Gene flow = (1-I_Sexual isolation_) × (1-I_PMPZ -isolation_)*
>
> *where I_Sexual isolation_ is index of sexual isolation and I_PMPZ isolation_ index of PMPZ isolation*

**Figure 11.**
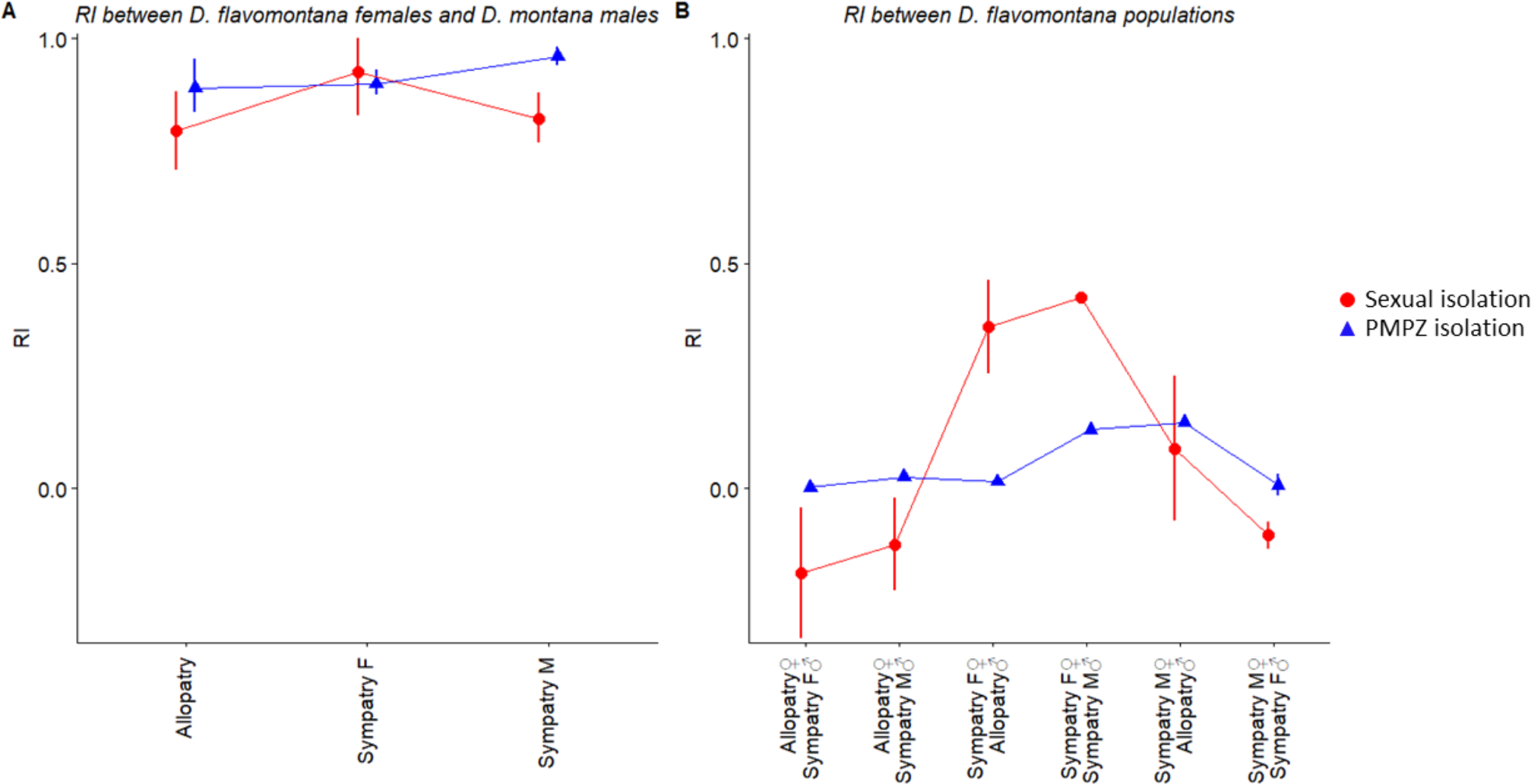
Reproductive isolation indices (RIs) calculated for sexual isolation and PMPZ isolation in different population types between *D. flavomontana* females and *D. montana* males and between populations of *D. flavomontana*. (A) In crosses between *D. flavomontana* females and *D. montana* males, sexual isolation was strongest in Sympatry F, while PMPZ isolation was strongest in Sympatry M. (B) Sexual and PMPZ isolation between *D. flavomontana* populations followed the same patterns as in interspecific matings: Sympatry F females discriminated against males of other populations (sexual isolation) and Sympatry M individuals showed lowered fertilization with some other conspecific population types (PMPZ isolation).

Given RI between *D. flavomontana* females and *D. montana* males, some gene flow may occur between species and this depends on population type. More gene flow is possible between individuals from Allopatric populations (0.027) compared to crosses in either Sympatric populations (Sympatry F = 0.005: Sympatry M = 0.009). A slight reduction in gene flow between *D. flavomontana* populations was evident between Sympatry F females and Allopatric males (0.632) or Sympatry M males (0.499), and between Sympatry M females and Allopatric males (0.776). Otherwise the probability for gene flow was 1 or higher.

## Discussion

Reinforcement of prezygotic reproductive barriers can enhance speciation both by strengthening reproductive isolation between sympatric species and by inducing divergence of reproductive traits between populations of the same species that live within and outside the area of sympatry (Howard 1993; Ortiz-Barrientos et al. 2009). A prerequisite for reinforcement is that postzygotic reproductive isolation is not complete, and producing interspecific hybrids is costly. In our study, postzygotic reproductive isolation between *D. montana* females and *D. flavomontana* males was nearly complete in all populations, leaving no chance for reinforcement to be currently acting. We also found in this cross nearly complete prezygotic isolation, potentially affected by reinforcement in the past. However, in crosses between *D. flavomontana* females and *D. montana* males both F_1_ hybrid viability and fertility was lower than in intraspecific crosses, providing the opportunity for reinforcement to occur. We subsequently found that the strength of sexual and PMPZ barriers varied between population types, as predicted for reinforcement. Yukilevich (2012) showed that concordant pre- and postzygotic isolation asymmetries in sympatry may have affected 60–83% of all sympatric Drosophila species and that they have enhanced premating isolation by 18–26%, on average. In crosses between *D. flavomontana* females and *D. montana* males, premating sexual isolation was 27% stronger in sympatric Rocky Mountains populations (Sympatry F) than in Allopatry, and PMPZ isolation was 25% stronger in sympatric western coast populations (Sympatry M) than in Allopatry.

We utilized a study system in which we could assess variation in patterns of reproductive isolation between populations with different sympatric histories and species abundancies compared with isolation between allopatric populations. Our first prediction was that in sympatric Rocky Mountains populations, where *D. flavomontana* is abundant (Sympatry F) and where the ancestral populations of both species are likely to be located (Stone et al. 1960), reinforcement is targeted on premating sexual isolation to minimize disadvantageous reproductive interactions with heterospecifics. Reinforcement of sexual isolation was predicted to increase female *D. flavomontana* discrimination against *D. montana* males, resulting in reproductive character displacement between and within the species in *D. flavomontana*’s key courtship cues and to generate sexual isolation between *D. flavomontana* populations (Fig. 1). Our data supported these predictions. In sympatric Rocky Mountains populations, reinforcement of sexual isolation between *D. flavomontana* females and *D. montana* males was detected as increased discrimination of *D. flavomontana* females against males of both *D. montana* and other *D. flavomontana* populations and as increased differences in *D. flavomontana*’s key courtship cues, CHC profiles, between the two species and among the sexes. This work contributes to other research demonstrating reinforcement targeting on sexual isolation between sympatric species (e.g. Noor 1995; Saetre et al. 1997; Rundle and Schluter 1998; Hoskin et al. 2005; Jaenike et al. 2006; Kronforst et al. 2007) and between populations of the same species (e.g. Lemmon 2009; Bewick and Dyer 2014; Kozak et al. 2015).

Our second prediction was that in sympatric western coast populations, where *D. flavomontana* is a new invader and still rare (Sympatry M), reinforcement can be targeted either on sexual or PMPZ barriers, depending on whether female *D. flavomontana* maintain high species-discrimination ability when rare (Fig. 1). Reinforcement was assumed to target sexual isolation and have similar consequences as in sympatric Rocky Mountains populations when female species-discrimination remains high even when females are from the rarer species. If, however, female mate recognition or preference has decreased due to the lack of conspecific mating partners, then reinforcement was predicted to be targeted on PMPZ barriers and to induce reproductive character displacement in traits maintaining these barriers both between the species and between *D. flavomontana* populations (Fig. 1). Results on sexual and PMPZ barriers in these populations fully support the second prediction; only PMPZ barriers, including the transfer/storage and/or use of heterospecific sperm in fertilization, showed signs of reinforcement in Sympatry M. Furthermore, *D. flavomontana* populations showed PMPZ barriers in the proportion of developing eggs in crosses between Sympatry M females and allopatric males and between Sympatry M males and Sympatry F females, indicating potential far-reaching effects of reinforcement also on *D. flavomontana* populations. Turissini et al. (2018) suggested that natural and sexual selection can drive the evolution of PMPZ barriers when reinforcement of sexual isolation is not possible, but our study is the first to demonstrate this.

One interesting difference between *D. montana* and *D. flavomontana* is that their species-recognition relies on different sensory modalities, with *D. montana* females using mainly auditory cues (male courtship song) and *D. flavomontana* females olfactory cues (CHCs). Female *D. montana* rarely accepts a male without hearing his courtship song, and can be fooled into mating with muted (wingless) heterospecific males by playing them conspecific song (Liimatainen et al. 1992; Saarikettu et al. 2005). *D. montana* song (mainly its carrier frequency and other sound pulse characters) also plays a major role in sexual selection (Ritchie et al. 1998) and both the song frequency and female preference for frequency varies between geographically isolated populations (Klappert et al. 2007). Thus, the variation we describe in male song frequency and the number of cycles in a sound pulse between *D. montana* populations with different evolutionary histories of overlap with *D. flavomontana* is likely to be due to sexual selection within the species rather than to reinforcement. For *D. flavomontana* females, CHCs were more important than song, but females mated even without receiving these cues through their antennal sense organs. Divergence between *D. flavomontana* populations and sexes in CHCs show clear signs of reinforcement at both species and population levels. While most of the cuticular substances were rather generic (occur in many insect species) and were found in most of our specimens, the methyl-branched alkanes made an interesting exception by being present only in *D. flavomontana*’s sympatric populations, and thus possibly playing an important role in the evolution of mate choice. Giglio and Dyer (2013) have detected shifts in male and female sensory modalities in the use of olfactory and gustatory cues between *Drosophila recens* and *Drosophila subquinaria*, as well as between *D. subquinaria* populations that are sympatric or allopatric with *D. recens*. They speculate that selection may have acted on sympatric *D. subquinaria* males to increase discrimination for species recognition by shifting the sensory cues females prefer for mating. Mate choice of *D. montana* and *D. flavomontana* from all population types relied on the same cues (auditory vs. olfactory), which suggests that the divergence in the use of these cues is not of recent origin.

PMPZ barriers occurring after mating, but before zygote formation, have only recently received attention as important suppressors of interspecific gene flow. Reinforcement of these barriers may, however, be a common and also quite rapid process (Castillo and Moyle 2014; Comeault et al. 2016; Turissini et al. 2018). For example, Matute (2010) detected an increase in PMPZ isolation between *D. yakuba* and *D. santomea* in sympatry, in which *D. yakuba* females depleted the sperm of *D. santomea* males faster than that of conspecific males. Here, we found PMPZ barriers were evident in both species in the receipt and/or storage of heterospecific sperm and in the effectiveness of the ejaculate in enhancing female egg laying and fertilization. Problems in sperm transfer and storage in the reciprocal crosses between *D. montana* and *D. flavomontana* could be partly due to species differences in female seminal receptacle length (*D. montana*: 3.43 mm; *D. flavomontana*: 10.54 mm) which positively correlates with sperm length (*D. montana*: 3.34 mm; *D. flavomontana*: 5.53 mm, Pitnick et al. 1999). Sperm storage, especially in matings between *D. montana* females and *D. flavomontana* males, may be diminished because the longer sperm of *D. flavomontana* may not properly interact with the shorter seminal receptacles of *D. montana*. If sperm are stored, reduced fertilization could be due to sperm not being able to be released from storage for fertilization and/or impairing penetration of the egg membrane after release (Howard 1999; Wirtz 1999; Lawniczak and Begun 2007; Howard et al. 2009; Kelleher et al. 2009). Reduction in fertilization was detected between *D. flavomontana* females and *D. montana* males, as well as between *D. flavomontana* from Sympatry M and other conspecific population types. PMPZ barriers have also been detected in other members of the *D. virilis* group, both between species (Sweigart 2010; Sagga and Civetta 2011; Ahmed-Braimah and McAllister 2012) and between *D. montana* populations from different geographical regions (Jennings et al. 2014b; Garlovsky and Snook 2018). Finally, ejaculate transfer to females may also induce an insemination reaction, in which a mass forms in the vagina that inhibits sperm storage and blocks oviposition (Patterson 1946; Knowles and Markow 2001). Insemination reactions have been detected in *D. montana* females after intraspecific matings (Wheeler 1947) and could be even more pronounced after mating with *D. flavomontana* males.

Several studies (e.g. Whitney et al. 2006; Abbott et al. 2013; Harrison and Larson 2014) have shown that incomplete reproductive barriers between sympatric species can lead to gradual incorporation of alleles from one species into the gene pool of a second one, which can enhance adaptation to new environmental conditions (Seehausen 2004; Currat et al. 2008; Abbott et al. 2013). Gene flow was estimated to be highest in crosses between allopatric *D. flavomontana* females and *D. montana* males, as predicted when reinforcement should not be acting, and reduced in these crosses from sympatric populations regardless of species abundance, as predicted when reinforcement is acting. Our next goal will be to quantify the magnitude and direction of gene flow between *D. montana* and *D. flavomontana* across these populations using whole-genome sequencing combined with studies on stress tolerances and habitat preferences. These studies will help to determine whether the species have received adaptive gene complexes from each other, and whether this could explain recent shifts in species distributions.

Ecological divergence is usually regarded as a prerequisite for the evolution of reproductive isolation through reinforcement (see Schluter 2009). The most notable ecological differences between *D. montana* and *D. flavomontana* are found in host-tree specialization and climatic adaptations (Patterson 1952). These differences could have played an important role during the first steps of speciation, giving the diverging species time to evolve reproductive barriers before being able to establish sympatry. *D. montana* populations on the western coast have adapted to live on lower altitudes, like *D. flavomontana*, and here the habitat differences between species are smaller than in the Rocky Mountains. Even though some of the populations may have been sympatric or allopatric for a shorter time than the others, comparisons between population types show clear signs of reinforcement. As Coyne and Orr (1997) state, the striking increase in prezygotic isolation seen in virtually all sympatric taxa suggests that the effect of sympatry is not only profound but also rapid.

In conclusion, reinforcement has been shown to play a key role both in enhancing and strengthening existing species boundaries, and the field of speciation is now beginning to evaluate not only the conditions under which reinforcement occurs but also its broader evolutionary and ecological consequences (Pfennig 2016). Speciation research also needs to consider the origin of barrier effects and the ways in which they are coupled, as strong barriers to gene flow will evolve only if multiple barrier effects coincide (Butlin and Smadja 2018). We show here that the target of reinforcement may change from sexual isolation to PMPZ barriers, if female discrimination towards heterospecific males decreases when the females are from the rarer species, and that the consequences of this change also can be detected between conspecific populations. Accordingly, we argue that the reliance of reproductive isolation on multiple barriers is also beneficial because the barriers can compensate each other in situations where reinforcement of some barriers is restricted.

## Acknowledgements

We would like to thank H. Järvinen for her help with the experiments, and people in the laboratory for the fly maintenance. We also thank E. Virtanen, A. Hiillos and E. Övermark for their contribution in Wolbachia studies. This work was supported by the grants from Academy of Finland (project 132619) and Ella and Georg Ehrnrooth Foundation to Anneli Hoikkala and Academy of Finland (projects 268214 and 272927) to Maaria Kankare.

## Appendix

**Figure A1.**
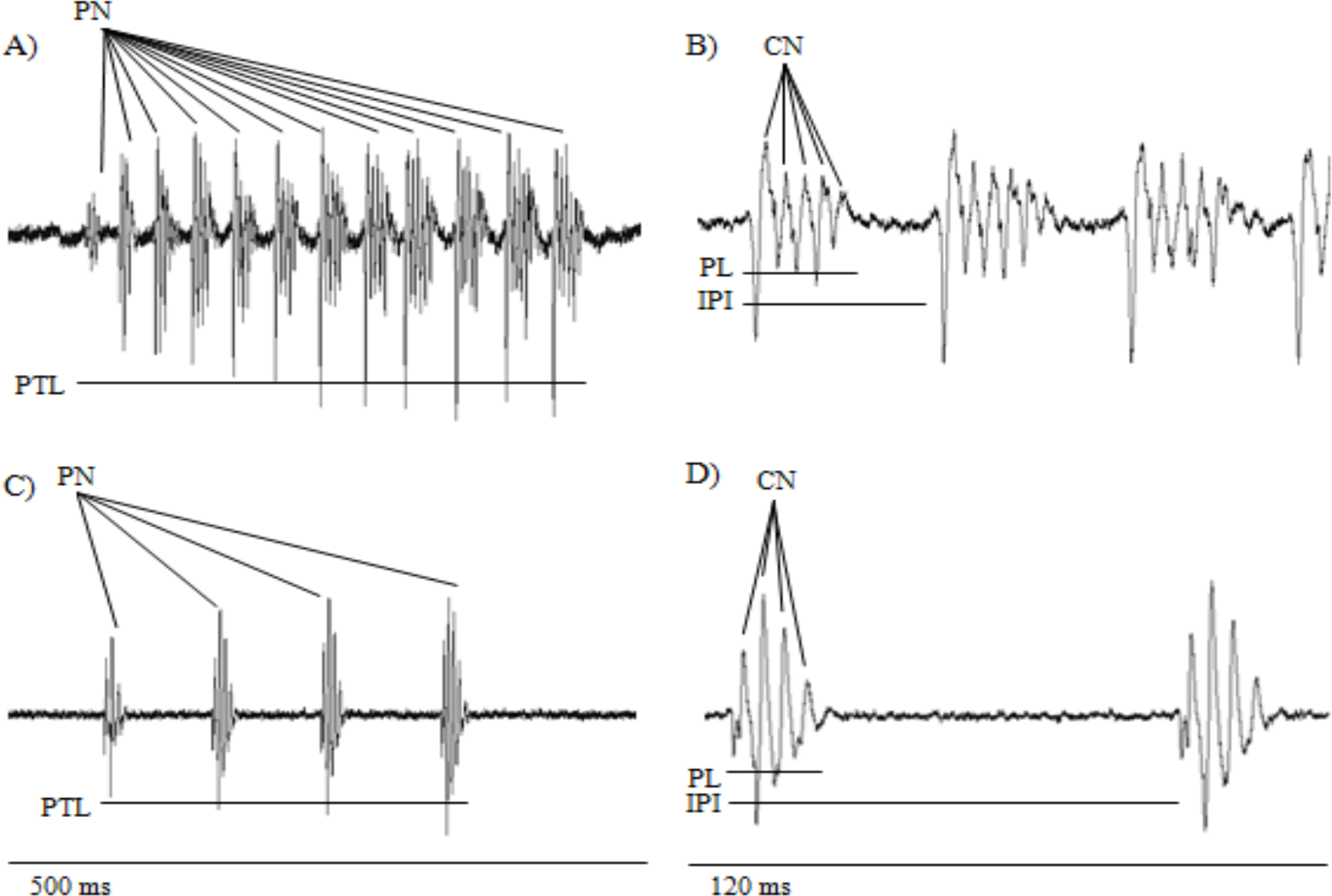
Oscillograms of the courtship songs of *D. montan*a (A, B) and *D. flavomontana* (C, D) males and the traits measured from them. PN = number of pulses in a pulse train, PTL = length of a pulse train, CN = number of cycles in a sound pulse, PL = length of a sound pulse, IPI = interpulse interval.

**Figure A2.**
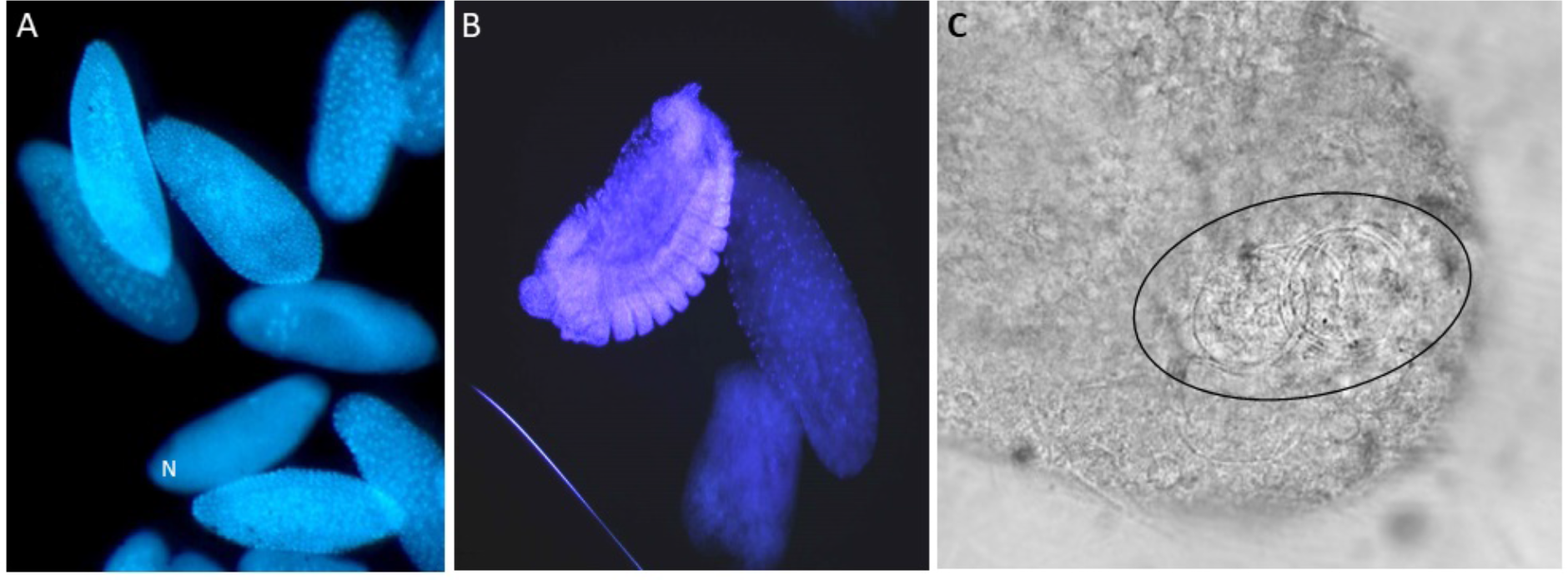
(A) Developing eggs with either clear mitotic division or (B) with cellular differentiation. Non-developing eggs had fewer than four nuclei visible within the egg (marked with N) (C) Sperm is visible as a spiral structure near the anterior end of the egg.

## References

Abbott, R., D. Albach, S. Ansell, J. W. Arntzen, S. J. E. Baird, N. Bierne, J. Boughman, A. Brelsford, C. A. Buerkle, R. Buggs, R. K. Butlin, U. Dieckmann, F. Eroukhmanoff, A. Grill, S. H. Cahan, J. S. Hermansen, G. Hewitt, A. G. Hudson, C. Jiggins, J. Jones, B. Keller, T. Marczewski, J. Mallet, P. Martinez-Rodriguez, M. Möst, S. Mullen, R. Nichols, A. W. Nolte, C. Parisod, K. Pfennig, A. M. Rice, M. G. Ritchie, B. Seifert, C. M. Smadja, R. Stelkens, J. M. Szymura, R. Väinölä, J. B. W. Wolf, and D. Zinner. 2013. Hybridization and speciation. J. Evol. Biol. 26:229–246.

Ahmed-Braimah, Y. H., and B. F. McAllister. 2012. Rapid evolution of assortative fertilization between recently allopatric species of *Drosophila*. Int. J. Evol. Biol. 1–9.

Arnold, M. L., and N. H. Martin. 2009. Adaptation by introgression. J. Biol. 8:82.

Bernier, U. R., D. A. Carlson, and C. J. Geden. 1998. Gas chromatography/mass spectrometry analysis of the cuticular hydrocarbons from parasitic wasps of the genus *Muscidifurax*. J. Am. Soc. Mass Spectrom. 9:320–332.

Bewick, E. R., and K. A. Dyer. 2014. Reinforcement shapes clines in female mate discrimination in *Drosophila subquinaria*. Evolution. 68:3082–3094.

Butlin, R. K., J. Galindo, and J. W. Grahame. 2008. Review. Sympatric, parapatric or allopatric: The most important way to classify speciation? Philos. Trans. R. Soc. B Biol. Sci. 363:2997–3007.

Butlin, R. K., and C. M. Smadja. 2018. Coupling, reinforcement, and speciation. Am. Nat. 191:155–172.

Carlson, J. R. 1996. Olfaction in Drosophila: From odor to behavior. Trends Genet. 12:175–180.

Carvajal-Rodriguez, A., and E. Rolan-Alvarez. 2006. JMATING: A software for the analysis of sexual selection and sexual isolation effects from mating frequency data. BMC Evol. Biol. 6:40.

Castillo, D. M., and L. C. Moyle. 2017. Conspecific sperm precedence is reinforced but sexual selection weakened in sympatric populations of *Drosophila*. bioRxiv 071886.

Castillo, D. M., and L. C. Moyle. 2014. Intraspecific sperm competition genes enforce post-mating species barriers in *Drosophila*. Proc. R. Soc. B Biol. Sci. 281:20142050.

Chenoweth, S. F., and M. W. Blows. 2006. Dissecting the complex genetic basis of mate choice. Nat. Rev.Genet. 7:681–692.

Christensen, R. H. B. 2018. Cumulative link models for ordinal regression with the R package ordinal. Submitted in J. Stat. Software.

Clark, N. L., J. E. Aagaard, and W. J. Swanson. 2006. Evolution of reproductive proteins from animals and plants. Reproduction 131:11–22.

Colyott, K., C. Odu, and J. M. Gleason. 2016. Dissection of signalling modalities and courtship timing reveals a novel signal in *Drosophila saltans* courtship. Anim. Behav. 120:93–101.

Comeault, A. A., A. Venkat, D. R. Matute. 2016. Correlated evolution of male and female reproductive traits drive a cascading effect of reinforcement in *Drosophila yakuba*. Proc. Biol. Sci. 283:23–48.

Coyne, J. A., and A. Orr. 1997. “Patterns of speciation in *Drosophila*” revisited. Evolution. 51:295–303.

Coyne, J. A., and A. H. Orr. 2004. Speciation. Sinauer Associates, Sunderland, MA.

Currat, M., M. Ruedi, R. J. Petit, and L. Excoffier. 2008. The hidden side of invasions: Massive introgression by local genes. Evolution. 62:1908–1920.

Dobzhansky, T. 1940. Speciation as a stage in evolutionary divergence. Am. Nat. 74:312–321.

Ferveur, J. F. 2005. Cuticular hydrocarbons: Their evolution and roles in *Drosophila* pheromonal communication. Behav. Genet. 35:279–295.

Garlovsky, M. D., and R. R. Snook. 2018. Persistent postmating, prezygotic reproductive isolation between populations. Ecol. Evol. 1–12.

Giglio, E. M., and K. A. Dyer. 2013. Divergence of premating behaviors in the closely related species *Drosophila subquinaria* and *D. recens*. Ecol. Evol. 3:365–374.

Gleason, J. M., A. A. Pierce, A. L. Vezeau, and S. F. Goodman. 2012. Different sensory modalities are required for successful courtship in two species of the *Drosophila willistoni* group. Anim. Behav. 83:217–227.

Harrison, R. G., and E. L. Larson. 2014. Hybridization, introgression, and the nature of species boundaries. J. Hered. 105:795–809.

Hoskin, C. J., and M. Higgie. 2010. Speciation via species interactions: The divergence of mating traits within species. Ecol. Lett. 13:409–420.

Hoskin, C. J., M. Higgie, K. R. McDonald, and C. Moritz. 2005. Reinforcement drives rapid allopatric speciation. Nature 437:1353–1356.

Howard, D. J. 1999. Conspecific sperm and pollen precedence and speciation. Annu. Rev. Ecol. Syst. 30:109–132.

Howard, D. J. 1993. Reinforcement: Origin, dynamics, and fate of an evolutionary hypothesis. Pp. 46–69 in R.G. Harrison, ed. Hybrid zones and the evolutionary process. Oxford University Press, NY.

Howard, D. J., S. R. Palumbi, L. M. Birge, and M. K. Manier. 2009. Sperm and speciation. Pp. 367–403 in T.R. Birkhead, D.J. Hosken and S. Pitnick, eds. Sperm Biology, an evolutionary persective. Academic Press.

Jaenike, J., K. A. Dyer, C. Cornish, and M. S. Minhas. 2006. Asymmetrical reinforcement and Wolbachia infection in Drosophila. PLoS Biol. 4:1852–1862.

Jennings, J. H., W. J. Etges, T. Schmitt, and A. Hoikkala. 2014a. Cuticular hydrocarbons of *Drosophila montana*: Geographic variation, sexual dimorphism and potential roles as pheromones. J. Insect Physiol. 61:16–24.

Jennings, J. H., R. R. Snook, and A. Hoikkala. 2014b. Reproductive isolation among allopatric *Drosophila montana* populations. Evolution. 68:3095–3108.

Kelleher, E. S., T. D. Watts, B. A. LaFlamme, P. A. Haynes, and T. A. Markow. 2009. Proteomic analysis of *Drosophila mojavensis* male accessory glands suggests novel classes of seminal fluid proteins. Insect Biochem. Mol. Biol. 39:366–371.

Klappert, K., D. Mazzi, A. Hoikkala, and M. G. Ritchie. 2007. Male courtship song and female preference variation between phylogeographically distinct populations of *Drosophila montana*. Evolution. 61:1481–1488.

Knowles, L. L., and T. A. Markow. 2001. Sexually antagonistic coevolution of a postmating-prezygotic reproductive character in desert *Drosophila*. Proc. Natl. Acad. Sci. 98:8692–8696.

Kozak, G. M., G. Roland, C. Rankhorn, A. Falater, E. Berdan, and R. Fuller. 2015. Behavioral isolation due to cascade reinforcement in *Lucania* killifish. Am. Nat. 185:491–506.

Kronforst, M. R., L. G. Young, and L. E. Gilbert. 2007. Reinforcement of mate preference among hybridizing *Heliconius* butterflies. J. Evol. Biol. 20:278–285.

Lawniczak, M. K. N., and D. J. Begun. 2007. Molecular population genetics of female-expressed mating-induced serine proteases in *Drosophila melanogaster*. Mol. Biol. Evol. 24:1944–1951.

Lemmon, E. M. 2009. Diversification of conspecific signals in sympatry: Geographic overlap drives multidimensional reproductive character displacement in frogs. Evolution. 63:1155–1170.

Liaw, A. and Wiener, M. 2002. Classification and regression by randomForest. R news. 2:18–22.

Liimatainen, J., A. Hoikkala, J. Aspi, and P. Welbergen. 1992. Courtship in *Drosophila montana*: the effects of male auditory signals on the behaviour of flies. Anim. Behav. 43:35–48.

Matute, D. R. 2010. Reinforcement of gametic isolation in *Drosophila*. PLoS Biol. 8:e1000341.

Matute, D. R. 2014. The magnitude of behavioral isolation is affected by characteristics of the mating community. Ecology and evolution. 4:2945–2956.

Mazzi, D., J. Kesäniemi, A. Hoikkala, and K. Klappert. 2009. Sexual conflict over the duration of copulation in *Drosophila montana*: Why is longer better? BMC Evol. Biol. 9:132.

Morales-hojas, R., M. Reis, C. P. Vieira, and J. Vieira. 2011. Molecular phylogenetics and evolution resolving the phylogenetic relationships and evolutionary history of the *Drosophila virilis* group using multilocus data. Mol. Phylogenet. Evol. 60:249–258.

Noor, M. A. 1995. Speciation driven by natural selection in *Drosophila*. Nature 375:674.

Noor, M. A. F. 1999. Reinforcement and other consequences of sympatry. Heredity. 83:503–508.

Nosil, P. 2012. Ecological Speciation. Oxford University Press, NY.

Nosil, P., L. J. Harmon, and O. Seehausen. 2009. Ecological explanations for (incomplete) speciation.Trends Ecol. Evol. 24:145–156.

Ortiz-Barrientos, D., A. Grealy, and P. Nosil. 2009. The genetics and ecology of reinforcement: Implications for the evolution of prezygotic isolation in sympatry and beyond. Ann. N. Y. Acad. Sci. 1168:156–182.

Päällysaho, S., J. Aspi, J. O. Liimatainen, and A. Hoikkala. 2003. Role of X chromosomal song genes in the evolution of species-specific courtship songs in *Drosophila virilis* group species. Behav. Genet. 33:25–32.

Patterson, J. 1946. A new type of isolating mechanism in *Drosophila*. Proc. Natl. Acad. Sci. 32:202–208.

Patterson, J. 1952. Revision of the montana complex of the virilis species group. Univerisity Texas Publ. 5204:20–34.

Pfennig, K. S. 2016. Reinforcement as an initiator of population divergence and speciation. Curr. Zool. 62:145–154.

Pinheiro J., Bates D., DebRoy S., Sarkar D. and R Core Team 2018. nlme: Linear and nonlinear mixed effects models. R package version 3.1-137, https://CRAN.R-project.org/package=nlme.

Pitnick, S., T. Markow, and G. S. Spicer. 1999. Evolution of multiple kinds of female sperm-storage organs in *Drosophila*. Evolution. 53:1804–1822.

Price, C. S. C., C. H. Kim, C. J. Gronlund, and J. A. Coyne. 2001. Cryptic reproductive isolation in the *Drosophila simulans* species complex. Evolution. 55:81–92.

R Core Team 2017. R: A Language and Environment for Statistical Computing. https://www.R-project.org/

Ritchie, M. G., R. M. Townhill, and A. Hoikkala. 1998. Female preference for fly song: Playback experiments confirm the targets of sexual selection. Anim. Behav. 56:713–717.

Rolan-Alvarez, E., and A. Caballero. 2000. Estimating sexual selection and sexual isolation effects from mating frequencies. Evolution. 54:30–36.

Rundle, H. D., and D. Schluter. 1998. Reinforcement of stickleback mate preferences: Sympatry breeds contempt. Evolution. 52:200–208.

Saarikettu, M., J. O. Liimatainen, and A. Hoikkala. 2005. The role of male courtship song in species recognition in *Drosophila montana*. Behav. Genet. 35:257–263.

Saetre, G. P., M. Král, and S. Bureš. 1997. Differential species recognition abilities of males and females in a flycatcher hybrid zone. J. Avian Biol. 28:259–263.

Sagga, N., and A. Civetta. 2011. Male-female interactions and the evolution of postmating prezygotic reproductive isolation among species of the *Virilis* subgroup. Int. J. Evol. Biol. 1–11.

Salminen, T. S., and A. Hoikkala. 2013. Effect of temperature on the duration of sensitive period and on the number of photoperiodic cycles required for the induction of reproductive diapause in *Drosophila montana*. J. Insect Physiol. 59:450–457.

Schluter, D. 2009. Evidence for ecological speciation and its alternative. Science. 323:737–741.

Seehausen, O. 2004. Hybridization and adaptive radiation. Trends Ecol. Evol. 19:198–207.

Servedio, M. R. 2001. Beyond reinforcement: The evolution of premating isolation by direct selection on preferences and postmating, prezygotic incompatibilities. Evolution. 55:1909–1920.

Servedio, M. R. 2009. The role of linkage disequilibrium in the evolution of premating isolation. Heredity. 102:51–56.

Servedio, M. R., and M. A. F. Noor. 2003. The role of reinforcement in speciation: Theory and data. Annu.Rev. Ecol. Evol. Syst. 34:339–364.

Simon, C., F. Frati, A. Beckenbach, B. Crespi, H. Lu, and P. Flook. 1994. Evolution, weighting, and phylogenetic utility of mitochondrial gene sequences and a compilation of conserved polymerase chain reaction primers. Ann. Entomol. Soc. Am. 87:651–701.

Skaug H., D. Fournier, A. Nielsen, A. Magnusson, B. Bolker. 2013. Generalized linear mixed models using AD model builder. R package version 0.7.

Smadja, C. M., and R. K. Butlin. 2011. A framework for comparing processes of speciation in the presence of gene flow. Mol. Ecol. 20:5123–5140.

Snook, R. R., and T. L. Karr. 1998. Only long sperm are fertilization-competent in six sperm-heteromorphic *Drosophila* species. Curr. Biol. 8:291–294.

Sobel, J. M., and G. F. Chen. 2014. Unification of methods for estimating the strength of reproductive isolation. Evolution. 68:1511–1522.

Stone, W. S., W. C. Guest, and F. D. Wilson. 1960. The evolutionary implications of the cytological polymorphism and phylogeny of the virilis group of *Drosophila*. Proc. Natl. Acad. Sci. 46:350–361.

Sweigart, A. L. 2010. The genetics of postmating, prezygotic reproductive isolation between *Drosophila virilis* and *D. americana*. Genetics 184:401–410.

Tauber, E., and D. F. Eberl. 2003. Acoustic communication in *Drosophila*. Behav. Processes 64:197–210.

The Marie Curie speciation network. 2012. What do we need to know about speciation? Trends Ecol. Evol. 27:27–39.

Throckmorton, L. H. 1982. The virilis species group. The Genetics and Biology of Drosophila. London Academy Press. 3:227–296.

Turissini, D. A., G. Liu, J. R. David, and D. R. Matute. 2015. The evolution of reproductive isolation in the *Drosophila yakuba* complex of species. J. Evol. Biol. 28:557–75.

Turissini, D. A., J. A. McGirr, S. S. Patel, J. R. David, and D. R. Matute. 2018. The rate of evolution of postmating-prezygotic reproductive isolation in *Drosophila*. Mol. Biol. Evol. 35:312–334.

Wheeler, M. R. 1947. The insemination reaction in intraspecific matings of *Drosophila*. Univerisity Texas Publ. 4720:78–115.

Whitney, K. D., R. A. Randell, and L. H. Rieseberg. 2006. Adaptive introgression of herbivore resistance traits in the weedy sunflower *Helianthus annuus*. Am. Nat. 167:794–807.

Wirtz, P. 1999. Mother species-father species: Unidirectional hybridization in animals with female choice.Anim. Behav. 58:1–12.

Wu, C. I., H. Hollocher, D. J. Begun, C. F. Aquadro, Y. Xu, and M. L. Wu. 1995. Sexual isolation in *Drosophila melanogaster*: a possible case of incipient speciation. Proc. Natl. Acad. Sci. U. S. A. 92:2519–2523.

Yukilevich, R. 2012. Asymmetrical patterns of speciation uniquely support reinforcement in *Drosophila*. Evolution. 66:1430–1446.

